# A mechanosensing mechanism mediated by IRSp53 controls plasma membrane shape homeostasis at the nanoscale

**DOI:** 10.1101/2021.08.01.454667

**Authors:** Xarxa Quiroga, Nikhil Walani, Albert Chavero, Alexandra Mittens, Andrea Disanza, Francesc Tebar, Xavier Trepat, Robert G. Parton, Giorgio Scita, Maria Isabel Geli, Marino Arroyo, Anabel-Lise Le Roux, Pere Roca-Cusachs

## Abstract

As cells migrate and experience forces from their surroundings, they constantly undergo mechanical deformations which reshape their plasma membrane (PM). To maintain homeostasis, cells need to detect and restore such changes, not only in terms of overall PM area and tension as previously described, but also in terms of local, nano-scale topography. Here we describe a novel phenomenon, by which cells sense and restore mechanically induced PM nano-scale deformations. We show that cell stretch and subsequent compression reshape the PM in a way that generates local membrane evaginations in the 100 nm scale. These evaginations are recognized by the I-BAR protein IRSp53, which triggers a burst of actin polymerization mediated by Rac1 and Arp2/3. The actin polymerization burst subsequently re-flattens the evagination, completing the mechanochemical feedback loop. Our results demonstrate a new mechanosensing mechanism for PM shape homeostasis, with potential applicability in different physiological scenarios.

**Teaser:** Cell stretch cycles generate PM evaginations of ≈100 nm which are sensed by IRSp53, triggering a local event of actin polymerization that flattens and recovers PM shape.

## Introduction

Cells constantly exchange information with their surroundings, and external inputs are first received by their outermost layer, the plasma membrane (PM). This interface, far from being an inert wall, integrates and transmits incoming stimuli, ultimately impacting cell behaviour. In this context, the traditional view of such stimuli as biochemical messengers has now changed to include the concept that physical perturbations are also of major importance (*1–3*). By sensing and responding to physical and biochemical stimuli, one of the main functions of the PM is to adapt to the changes in shape that cells experience as they migrate or are mechanically deformed, in a variety of physiological conditions (*4–9*). To date, research in this area has largely focused on the regulation of PM area and tension, at the level of the whole cell (*10–12*). For instance, cell stretch or decrease in medium osmolarity have been commonly used to raise PM tension, unfolding membrane reserves (ruffles, caveolae), inhibiting endocytosis and promoting exocytosis (*13–17*). Conversely, cell exposure to a hypertonic solution or cell compression have been employed to decrease PM tension, leading to an increase on the activity of different endocytic pathways (*18–21*). These studies have shown that PM tension homeostasis is maintained by regulating PM area through mechanisms like endocytosis, exocytosis, or the assembly and disassembly of PM structures like ruffles and caveolae.

However, changes in cell PM area upon mechanical perturbations are necessarily accompanied by changes in topography at the local scale. This is clearly exemplified by caveolae flattening upon cell stretch (*22*) or creation of PM folds at the sub-μm scale upon cell compression (*20*). Curvature also arise when membranes are exposed to either external topographical cues (*23, 24*) or internal pulling by actin filaments (*25–27*). To maintain PM homeostasis, cells should thus be able to not only respond to overall changes in PM tension or area, but also to local changes in PM topography. This requirement is even clearer if one considers recent findings showing that tension does not propagate extensively throughout the whole ensemble of the PM, but dissipates in small areas of less than 5 μm (*28*). However, if such local PM shape homeostasis mechanisms exist, and how they operate, is still unknown.

Here, we studied this problem by using as a model the controlled compression of fibroblasts through the application and release of stretch. We show that upon cell compression, bud-shaped PM deformations of negative curvature (evaginations) on the 100 nm scale are formed and enriched by IRSp53, a negative curvature-sensing protein. This creates a local node where specific PM topography is selectively coupled through IRSp53 to activate actin polymerization mediated by Rac1 and Arp2/3. The activation of this cascade flattens the structure, recovering the PM shape to its initial state. Our findings demonstrate a local mechanosensing mechanism that controls PM homeostasis when perturbed through compression.

## Results

### Compression generates dynamic PM evaginations of 100 nm in width

To study how PM topography is regulated, we submitted normal human dermal fibroblasts (NHDF) transfected with an EGFP-membrane marker to a physiologically relevant 5% equibiaxial stretch by using a custom-made stretch system composed by a PDMS stretchable membrane clamped between two metal rings, as previously described in (29) (see methods). Cell response during and after stretch was monitored by live fluorescence imaging. As previously described, when tensile stress was applied cells increased their area by depleting PM reservoirs, such as ruffles (10, 20). After 3 minutes, stretch was released, resulting in a compression stimulus. At this point, excess membrane was stored again in folds, visualized as bright fluorescent spots of ≈ 500 nm (Fig. 1A and Supp. Video SV01). These spots incorporate approximately 1.5% of PM area (Supp. Fig. 1A), and thus store an important fraction of the area modified by cell stretch. As we have previously published (20), these spots are formed passively by the PM to accommodate compression, analogously to what occurs when compressing synthetic lipid bilayers (30). In cells however, passive fold formation is followed by active resorption involving actin cytoskeleton rearrangements, allowing for topography equilibration within 2 minutes (Fig. 1B). As the diffraction limit of a standard fluorescence microscope lays in the range of 500 nm, we characterized the structure of the compression-generated folds in more detail using electron microscopy. Cells transfected with a PM marker were seeded in a 3D patterned PDMS membrane, stretched and immediately fixed after the release of the stimulus. Next, brightfield and fluorescent images of the 3D pattern and the cells on it were acquired and samples were further processed for SEM imaging. Computational alignment tools allowed for correlation between brightfield, fluorescence and SEM images. De-stretched cells displayed numerous bud-shaped evaginations at their apical PM side that correlated with the bright spots seen by fluorescence (Figs. 1C and D), showing that the PM bends outwards (thereby minimizing friction with the underlying cortex). To accurately measure the size of these evaginations we moved into transmission electron microscopy (TEM). By comparing non stretched to stretched-released cells, we observed that the first displayed a homogeneously flat PM, while the second group displayed bud-shaped evaginations on the apical side (Fig. 1E). Analysis of the shape profile of compression-induced evaginations yielded an average diameter in the neck (cylindrical shape) of 83 nm and of 115 nm in the head (spherical shape), and average curvatures of 0.03 and 0.02 nm^-1^ respectively (Figs. 1F, G and H). These data indicate that PM compression led primarily to the formation of evaginations of regular size and shape at the apical side, which are immediately resorbed by the cell in an active process to re-equilibrate PM topography and tension.

**Fig. 1.**
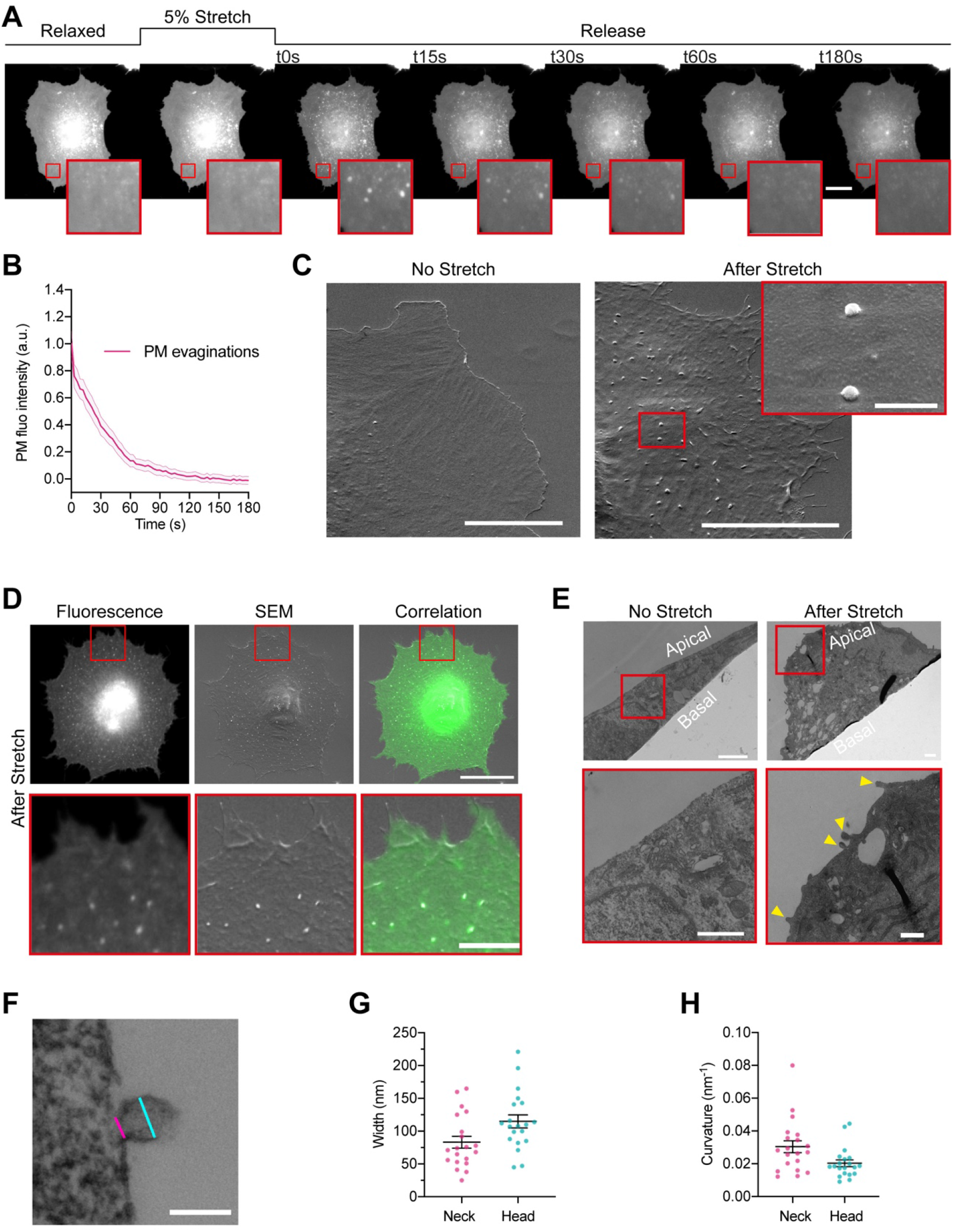
Cellular stretch generates PM evaginations with a defined curvature. **(A)** Time course images of a NHDF transfected with EGFP-membrane marker before, during and after 5 % constant stretch application. PM evaginations are seen as bright fluorescent spots after the release of the stretch due to compression of the PM. Scale bar is 20 μm. **(B)** Dynamics of PM evaginations after stretch release quantified as the change in fluorescence of the structure with time. N=12 cells from 3 independent experiments. **(C)** NHDF imaged through SEM. A non-stretched cell (left), and a cell just after stretch release (right) are shown. Scale bars are 10 μm in main images, 500 nm in magnified image (framed in red). **(D)** Correlation between fluorescence and SEM images of a non-stretched and stretched-released NHDF. Matching was achieved by using a patterned substrate together with computational tools for alignment. Scale bar is 20 μm for the main images and 2 μm for the insets. **(E)** TEM images of a non-stretched and a stretched-released NHDF. Yellow arrows in magnified image point at PM evaginations formed at the apical side of the cell. Scale bars are 1μm for the main images and 500 nm for the insets. **(F)** Detail of an evagination, cyan and magenta lines show evagination’s head and neck diameters, respectively. Scale bar is 100 nm. **(G, H)** Corresponding evagination neck and head diameters **(G)** and curvatures **(H)**. N=22 evaginations from 3 independent experiments. Data show mean ± s.e.m. In A, C, D, and E, red-framed images show a magnification of the areas marked in red in the main image.

### Actin is recruited to evaginations through the curvature-sensing protein IRSp53

In light of these results, we wondered if the PM evaginations formed upon compression could be detected by the cell, triggering a mechanism to recover PM shape. Based on previous results showing that actin depolymerization by either Latrunculin A or Cytochalasin D blocked PM remodeling after stretch (20), we hypothesized that the first step for recovery likely involved reattachment of the evaginated PM to the actin cortex. To explore this idea, we submitted NHDFs to a cycle of stretch and we imaged their response after stretch release. To visualize actin dynamics, cells were co-transfected with a PM marker together with a plasmid expressing an actin nanobody bound to a GFP fluorophore (ACG). As evaginations were being resorbed, actin was recruited to the same spot (Fig. 2A and Supp. Video SV02). Quantification of fluorescence intensity of PM and ACG markers showed a recruitment of actin which was delayed with respect to the PM marker (Figs. 2B and C), reaching a maximum at 15 s. This was followed by a decrease in the intensity of both markers that concluded when evaginations were resorbed (Fig. 2B). This suggests that the PM quickly reattaches to the underlying cortex, which then mediates remodeling of the structure. To further confirm the hypothesis and to prevent any mechanical interference caused by actin manipulation (31), we repeated the same experiment over-expressing the PM-cortex linker ezrin (32, 33). mEmerald-Ezrin also co-localized with evaginations during their resorption (Fig. 2D and Supp. Video SV03) and fluorescence analysis of PM and ezrin markers revealed a recruitment of the protein that mimicked, with a delay of 10 s, the one seen with actin (Figs. 2E and F).

**Fig. 2.**
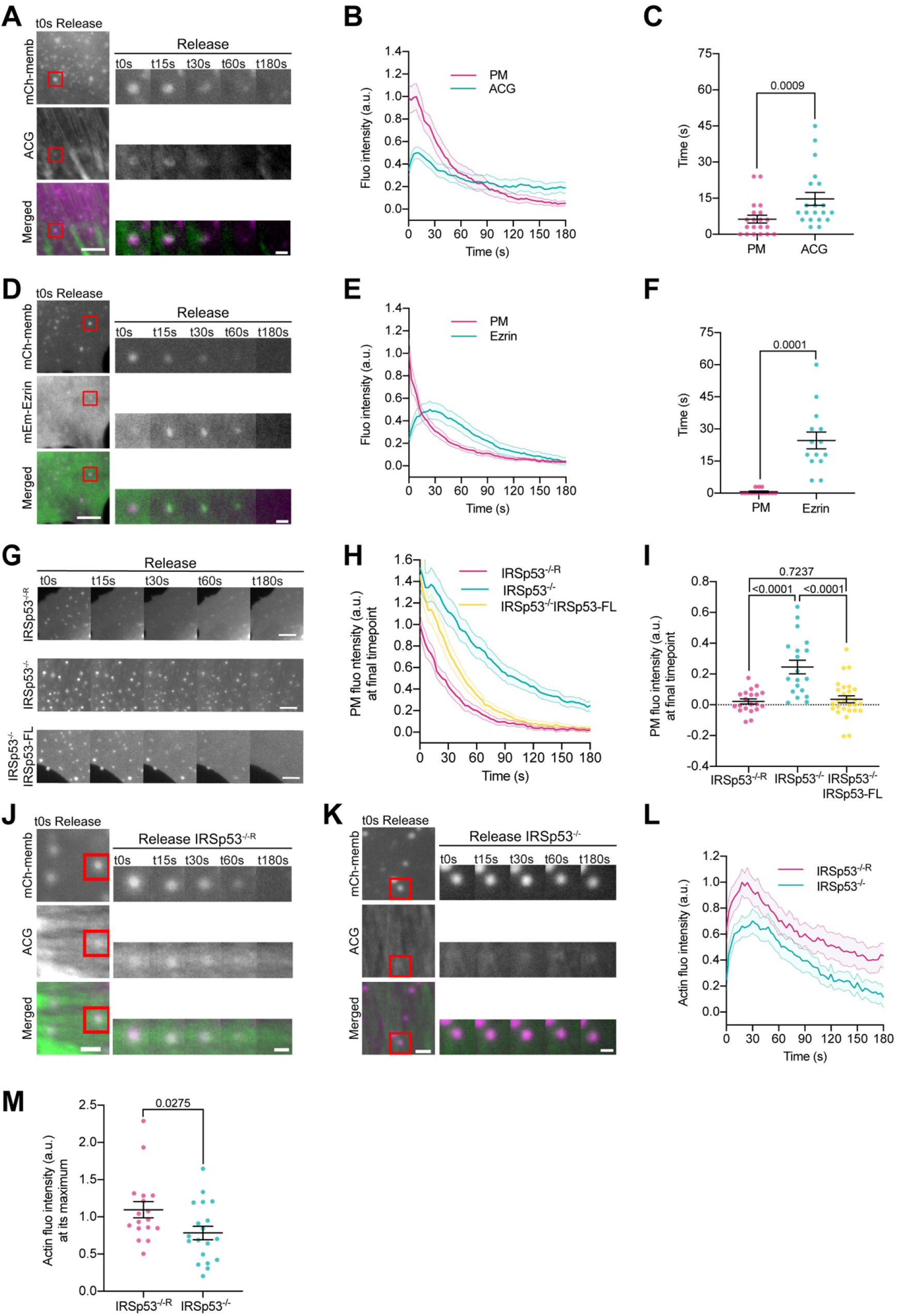
PM evaginations trigger local actin recruitment mediated by the IBAR protein IRSp53. (**A**) Time course images of mCherry-membrane and Actin Chromobody-GFP (ACG) marking PM evaginations in NHDF after stretch release. **(B)** Dynamics of PM evaginations quantified through mCh-membrane or ACG fluorescence markers during stretch release in NHDF. N= 20 cells from 3 independent experiments. **(C)** Timepoint of maximal fluorescence intensity of PM and ACG. Statistical significance was assessed through Wilcoxon test. N= 20 cells from 3 independent experiments. **(D)** Time course images of mCherry-membrane and mEmerald-Ezrin marking PM evaginations in NHDF after the release of the stretch. **(E)** Dynamics of PM evaginations quantified through mCh-membrane and mEmerald-Ezrin fluorescence markers after stretch release in NHDF. N= 14 cells from 2 independent experiments. **(F)** Timepoint of maximal fluorescence intensity of PM and ezrin markers. Statistical significance was assessed through Wilcoxon test. N= 14 cells from 2 independent experiments. **(G)** Time course images of PM evaginations tagged by mCherry-membrane in IRSp53^-/-R^, IRSp53^-/-^ and IRSp53^-/-^ EGFP-FL-IRSp53 cells after the release of stretch. **(H)** Dynamics of PM evaginations (mCh-membrane marker) after stretch in IRSp53^-/-R^, IRSp53^-/-^ and IRSp53^-/-^ EGFP-FL-IRSp53. **(I)** Differences in PM fluorescence intensity at the final timepoint of acquisition (180s after the release of the stretch). Significant differences were tested through ANOVA. N= 20, 19 and 28 cells from 3, 3 and 5 independent experiments. **(J, K)** Time course images of mCherry-membrane and ACG marking the evolution of both PM evaginations and actin after the release of the stretch in IRSp53^-/-R^ MEF cells (J) and IRSp53^-/-^ MEF (K). **(L)** ACG dynamics at PM evaginations after stretch in both IRSp53^-/-R^ and IRSp53^-/-^ MEF. **(M)** Maximal fluorescence intensity of ACG during the resorption process for IRSp53^-/-R^ and IRSp53^-/-^ cells. Statistical significance was assessed through Man-Whitney test. N= 17 and 19 cells from 4 independent experiments. For panels A, D, and G, scale bars are 5 μm and 1 μm for insets. For panels J and K, scale bars are 2 μm and 1 μm for insets. Data show mean ± s.e.m.

The burst in actin polymerization at the evaginated PM and the simultaneous reattachment to the cortex suggest that the local topography generated by compression may act as the mechanical input triggering the subsequent polymerization event. Indeed, membrane curvature can recruit different signaling molecules (19, 34–37), chief among them curvature-sensing BAR proteins (38–41). The superfamily of BAR proteins includes molecules containing different curvature sensing and generating BAR domains: The N-BAR and F-BAR domains, which interact with positively curved membranes (invaginations), and the I-BAR domain for the opposite type of curvature (negatively curved membranes or evaginations). In addition, these proteins also contain other domains, many of them reported to recruit actin nucleation promoting factors (NPFs) or even directly binding actin monomers (42). Interestingly, a recent work described how ezrin needs to act in partnership with the I-BAR protein IRSp53 to enrich in negatively curved membranes (43). Previous work done on IRSp53 has related this protein to PM ruffling (44, 45), filopodia formation (46–49) and endocytosis (50), but, so far, no mechanosensing mechanism relying on its capacity to bind negatively-curved membranes has been described. Moreover, recent studies in vitro and in vivo have pointed out that the I-BAR domain of IRSp53 displays a peak of sorting at evaginations with curvatures of 0.05 nm^-1^, and that lower curvature values comparable to the ones obtained by TEM imaging of our evaginations also led to a two-fold enrichment of this domain with respect to a control membrane marker (47, 51).

Prompted by this idea, we tested if IRSp53 could be the molecular linker between PM shape and actin dynamics in our system. To do so, we created stable cell lines expressing IRSp53 shRNA and control Non-Targeting shRNA (NT-shRNA). By plotting the decrease in PM fluorescence at the location of the evagination as a function of time for both control and IRSp53 silencing, we compared how lack of this protein affected the resorption process of PM evaginations (Supp. Figs. 1B and C and Supp. Video SV04). To assess the effectiveness of resorption (and since not all curves in all conditions could be fitted to an exponential equation with a characteristic time scale), we compared residual PM fluorescence at the end of experiments, 180 s. Full reabsorption of evaginations leads to a complete return to fluorescent baseline, while presence of a residual fluorescence indicates non-reabsorbed evaginations. Concordant with our hypothesis, IRSp53-depleted NHDFs did not complete evaginations resorption after 180 s (Supp. Figs. 1C, D and Suppl. Video SV04), even though they stretched by the same amount as non-depleted cells (Supp. Fig. 1E and Supp. Video SV05). To corroborate this result, we used isogenic mouse embryonic fibroblasts (MEFs) derived from IRSp53 null mice, that were stably infected either with a control (IRSp53^-/-^) or an IRSp53-retroviral vector (IRSp53^-/-R^) to restore expression levels of IRSp53 similar to wild type fibroblasts, as previously described (52-54). IRSp53^-/-^ cells also stretched by the same amount as IRSp53^-/-R^ cells (Supp. Fig. 1F) and did not display significant changes in the number of evaginations generated after compression (Supp. Fig. 1G) or in the area stored by those (Supp. Fig 1H). However, and reinforcing the previous result, they showed a severe impairment in the resorption of the evaginations even 180 s after stretch release. We further re-introduced EGFP-tagged full-length (FL) wild type IRSp53 into IRSp53^-/-^ by transient transfection. The reintroduction of IRSp53-FL rescued the phenotype, leading to a full recovery of PM topography by resorbing the compression-generated evaginations in a lapse of 90 s (Figs. 2G-I and Supp. Videos SV06, SV07, SV08). Further, IRSp53^-/-^ cells exhibited a recruitment of actin to PM evaginations that was weakened with respect to IRSp53^-/-R^ cells (Fig. 2J-M and Supp. Videos SV09 and SV10), illustrating that actin assembly at the PM evaginations is dependent on the presence of the I-BAR protein.

Next, we tested whether the effect of IRSp53 in PM reshaping was local at evaginations, or a general non-specific cell-level effect due to the ability of IRSp53 to organize different NPFs (55, 56). To this end, we generated PM folds of very different nature and curvature. We transiently exposed cells to hypo-osmotic medium, leading to cell swelling. As previously described, re-exposure to iso-osmotic medium generates a water outflow from cells. For cells seeded on non-porous substrate such as PDMS, expelled water becomes trapped between the cell and the substrate, forming the dome-shaped invaginations known as vacuole-like dilations (VLDs). VLDs are much larger than compression-generated bud-shaped evaginations (several μm in size), with much lower curvature, and resorb in the order of minutes (20). Confirming the local, evagination-specific effect of IRSp53, VLD resorption was equivalent in IRSp53^-/-R^ and IRSp53^-/-^ cells (Supp. Fig. 2A, B, C and Supp. Videos SV11 and SV12). IRSp53 has also been related to actin polymerization in lamellipodia (44, 57). To discard that flattening of the evaginations was due to potential lamellipodial extension (cell spreading) after compression, we analyzed cell spreading dynamics. After a stretch-release cycle, cells did extend lamellipodia and spread during approximately 1 minute (Suppl. Fig. 2D). However, the time constant of spreading (obtained by fitting an exponential curve to the experimental curve) and the amount of area recovered were not altered by the loss of IRSp53 (Supp.Fig. 2E-G), discarding a role of this process in the resorption of evaginations.

### Homeostasis recovery after stretch requires integrity of SH3 and IBAR IRSp53 domains

So far, we have shown that PM remodeling of compression-generated evaginations is a local event, which depends on IRSp53 to organize a burst of actin polymerization that flattens the PM. Next, we investigated if this could be part of a mechanosensing mechanism. Indeed, the I-BAR domain of IRSp53 may recognize the curvature generated at the evaginations and further recruit NPFs to coordinate the polymerization event. However, IRSp53 possesses multiple domains with multiple interactors, as illustrated in Fig. 3A. First, the I-BAR domain of IRSp53 can not only interact with charged curved membranes, but also possesses a Rac Binding domain (RCB) which enables binding to activated Rac. Additionally, it has been described to bundle actin (58). IRSp53 also contains an atypical CRIB domain that binds to activated Cdc42, but not Rac1 (59) and, further, an SH3 domain that recruits different NPFs, such as WAVE2, Eps8 or N-WASP (55). To test the role of these different domains, we used a cohort of IRSp53 mutants each affecting a specific domain and impeding a specific interaction, as described in Fig. 3B. EGFP-labelled mutants disrupting the function of IBAR, CRIB and SH3 domains were expressed in the background of IRSp53^-/-^ cells, and PM remodeling after stretch was analyzed. Whereas a set of mutants was able to rescue the wild type phenotype (Figs. 3C, D and F), another group was not (Fig. 3C, E and G). The I-BAR mutant 4KE, in which positively charged Lysines 142, 143, 145 and 147 belonging to a basic patch involved in PM and actin binding were replaced by negatively charged Glutamic Acid to disrupt this interaction (58, 60), rescued the phenotype (Supp. Video SV13). Mutation of these amino acids was probably not efficient enough in preventing PM binding. Phenotype recovery was also observed with the I268N mutant in the CRIB domain, which impairs the interaction with Cdc42 (Supp. Video SV14). However, the full deletion of the I-BAR domain or point mutations I403P and W413G in the SH3, that impair the association of IRSp53 with all its SH3 interactors, including WAVE2 (61), VASP and Eps8 (52, 62), did not rescue homeostasis recovery after stretch release (Supp. Videos SV15, 16 and 17). Moreover, the over-expression of the I-BAR domain alone also failed to rescue the phenotype (Supp. Video SV18), suggesting that the interaction with the PM and active Rac1 is not sufficient to drive PM flattening in response to stretch. This ensemble of results points at a mechanism where the I-BAR domain of IRSp53 would interact with the curved membrane of evaginations, leading to actin polymerization via active Rac1 and activation of NPFs through its SH3 domain.

**Fig. 3:**
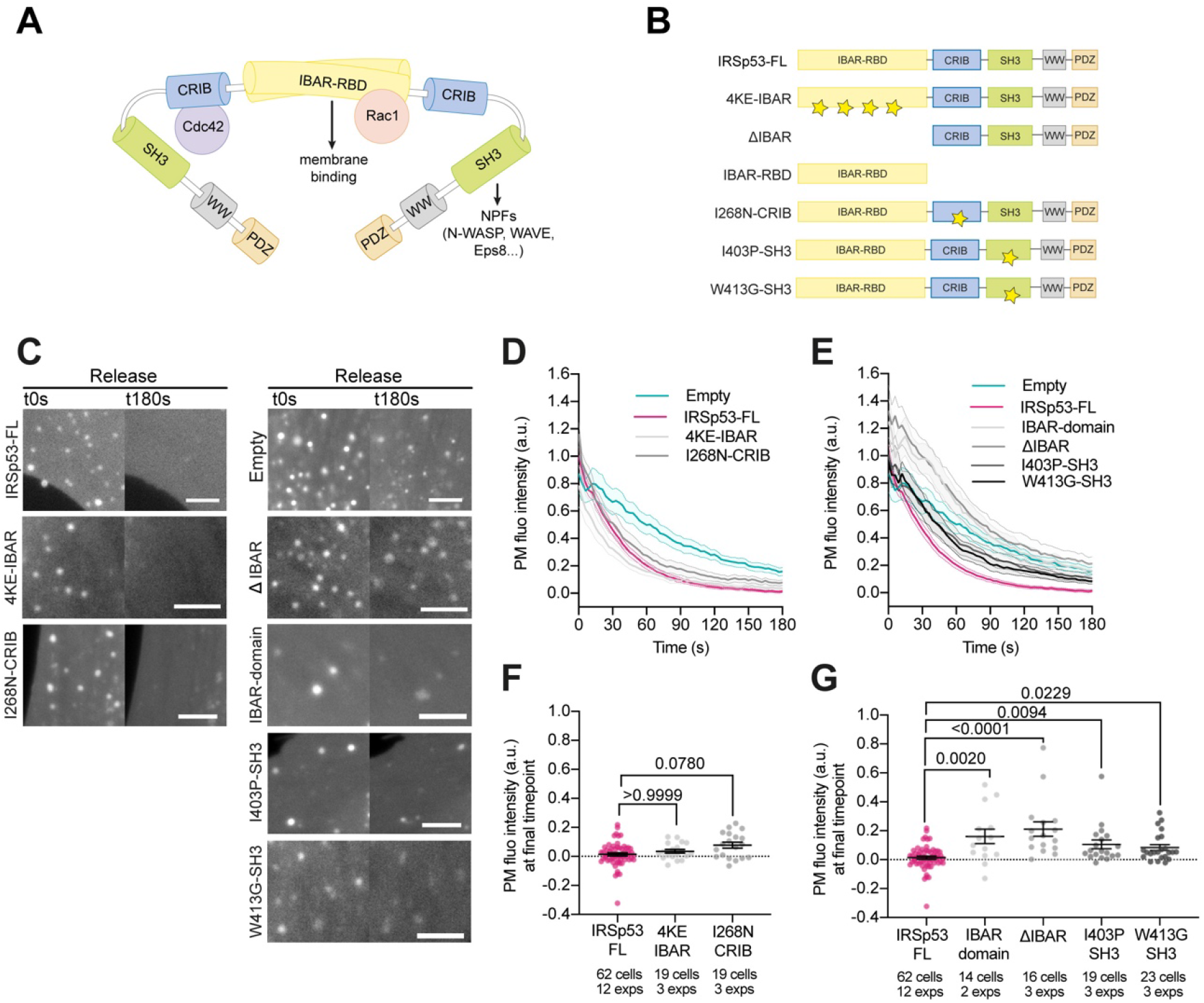
IBAR and SH3 domains of IRSp53 regulate the resorption of PM evaginations. **(A)** Schematics representing the IBAR protein IRSp53 and the different molecules interacting with its different domains. **(B)** Schematics of the IRSp53 mutants used in this study. Stars denote the location of mutations impairing the function of the different domains. **(C)** Images of PM evaginations of IRSp53^-/-^ cells transfected with mCh-membrane alone (empty) or in combination with the different full length or mutant forms of EGFP-IRSp53 at the first (t0 s) and last (t180 s) timepoint of acquisition after stretch. Scale bars are 5 μm. **(D-E)** Time course dynamics of PM evaginations of mCh-membrane transfected IRSp53^-/-^ cells either empty or reconstituted with the different full length or mutant forms of IRSp53. (D) shows IRSp53 mutants that rescue PM recovery after stretch, (E) shows IRSp53 mutants that do not rescue PM recovery after stretch. **(F-G)** Corresponding fluorescence intensity of PM evaginations at the last timepoint of acquisition (t180 s) after the release of stretch. Statistical significance was assessed through Kruskal-Wallis test. Data show mean ± s.e.m.

### IRSp53 acts as a mechanosensor by recognizing mechanically-induced PM curvature

To evaluate whether IRSp53 itself was directly recruited to evaginations, we imaged the dynamics of the EGFP-IRSp53-FL or mutant forms, expressed in IRSp53^-/-^ cells. Colocalization of the fluorescently labeled protein, either WT or mutated, and the PM marker was found in all cases (Supp. Fig. 3A-G), indicating that the presence of IRSp53 in the PM is not mediated exclusively by the I-BAR domain and rather occurs as an interplay of all different domains, as already suggested in previous studies (63, 64). We next analyzed the dynamics of EGFP-IRSp53-FL at the resorbing evaginations. Because IRSp53 is already bound to the PM, colocalization of the protein with the evaginations was observed from the first timepoint after stretch release. However, the decay in fluorescence of the IRSp53 coupled fluorophore was significantly slower than that of the PM marker (Figs. 4C and D), indicating that there is a progressive enrichment of IRSp53 to the evaginations while those are disappearing. To confirm this, we used APEX technology (65, 66) to visualize IRSp53 at PM evaginations using TEM. We co-transfected IRSp53^-/-^ cells with csAPEX2-GBP, a conditionally stable APEX marker bound to a nanobody specifically recognizing GFP (also called GFP-binding protein, GBP), and either EGFP-IRSp53-FL or a GFP-bound mitochondrial marker (Mito-GFP). As expected, a strong APEX signal (visible as a darker signal in the TEM image) was observed around the mitochondrial membrane for Mito-GFP-transfected cells (Supp. Fig. 3M), and at the tip of filipodia for EGFP-IRSp53-FL transfected cells (Supp. Fig. 3N) (47, 50). Confirming that IRSp53 is recruited to PM evaginations generated by a stretch-release cycle, such evaginations showed an increase in APEX signal in IRSp53-FL transfected cells (Fig. 4E), but not in control mito-GFP transfected cells (Fig. 4F).

**Fig. 4:**
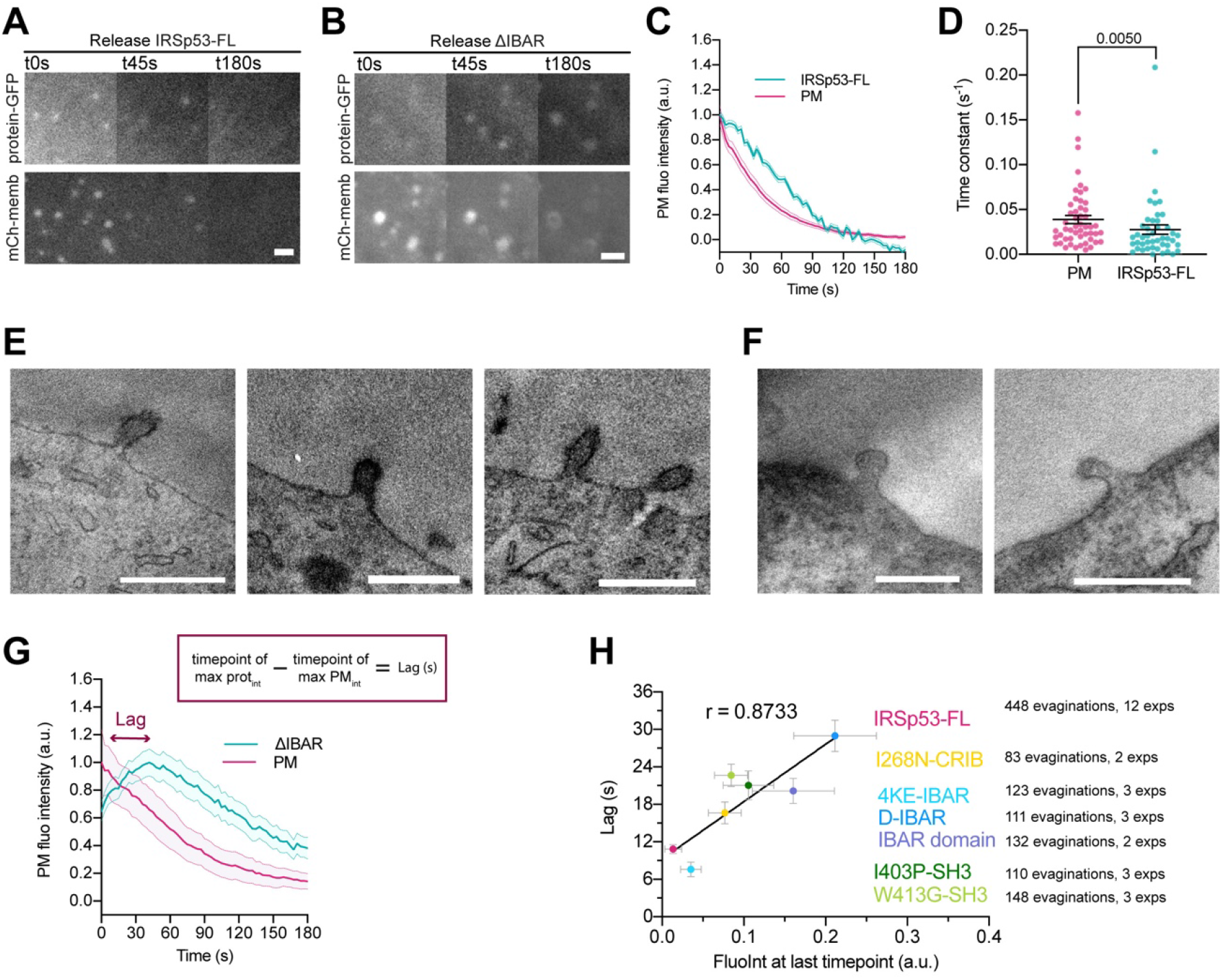
IRSp53 acts as a mechanosensor of PM curvature. **(A, B)** Images after stretch release of IRSp53^-/-^ cells transfected with mCh-membrane and either FL or ΔIBAR forms of IRSp53 coupled to EGFP. Scale bars are 2μm. **(C)** Dynamics of PM evaginations upon stretch release quantified through mCh-membrane or EGFP-IRSp53-FL fluorescence. **(D)** Time constants obtained by exponential fitting of the evagination resorption curves in the PM and EGFP-FL-IRSp53 channels. Statistical significance was assessed through Mann-Whitney test. N=53 cells from 12 independent experiments. **(E-F)** TEM images of PM evaginations coming from cells co-transfected with either APEX-GBP and (E) EGFP-IRSP53-FL or (F) control condition mito-GFP. APEX staining can be observed at the PM evaginations of EGFP-IRSp53-FL transfected cells marking IRSp53 position. Scale bars are 500 nm. **(G)** Dynamics of PM evaginations upon stretch release quantified through mCh-membrane or EGFP-ΔIBAR fluorescence. The purple arrow indicates the lag between the PM and IRSp53 signals, i.e., the time difference between the peaks of maximum intensity of both markers. N=12 cells from 3 independent experiments. **(H)** Time lag of FL or mutated IRSp53 plotted against the intensity of fluorescence at the last timepoint of acquisition. R indicates the Pearson correlation coefficient between both variables. Data show mean ± s.e.m.

Given that IRSp53 recognizes evaginations, we checked whether different mutants recognized the structure differently. The fluorescence dynamics of the IRSp53 mutants that had little effect on PM evaginations resorption followed a similar decay to the WT form (Supp. Figs. 3H and I). However, the mutants slowing down the resorption followed different dynamics, with an initial recruitment phase before the decay in fluorescence (Fig. 4G and Supp. Figs. 3J-L). This is illustrated in its most prominent example by the ΔIBAR mutant (Fig. 4G). These results indicate a delay in the recruitment of IRSp53 WT to the curved PM upon stretch as well. Indeed, due to the experimental time required to refocus samples and start imaging after compression (around 5-10 s), our time lapses fail to capture the process of PM evagination formation, or the recruitment of WT IRSp53. However, when the process is impaired due to IRSp53 mutations, dynamics are slowed down and we can capture the recruitment phase.

This led us to hypothesize that, although presence at the PM is a feature that does not depend on a single domain, recruitment at the curved evaginations could define the efficiency of homeostasis recovery after stretch. To quantify this, we measured the lag time between the timepoints of maximum intensity of the fluorescence signals of the PM and of the different IRSp53 mutant proteins (as illustrated in Fig. 4G). Confirming our hypothesis, plotting the lag time against the PM fluorescence intensity of the PM marker after 180 s (used previously as a marker for the efficiency of resorption of the evaginations) led to a strong positive correlation (Fig. 4H): the more IRSp53 recruitment was delayed with respect to the PM marker, the less efficient the resorption was. Removal of the I-BAR domain displayed the longest lags and the least efficient resorption, supporting the idea that curvature sensing through this domain is needed to couple the mechanical stress to active PM remodeling. If the domain is absent, IRSp53 cannot perform a quick binding to the evagination and start the mechanochemical loop. I403P and W413G mutations of the SH3 domain also led to long lags and inefficient resorption. Although in this case the I-BAR domain is not impaired, lack of interaction of IRSp53 with NPFs, which could be already bound to the PM and target IRSp53 there (56, 64), could delay both recruitment and the subsequent resorption process. Similarly, the I268N-CRIB mutant is probably delayed due to the lack of interaction with active Cdc42 already bound to the PM. Nevertheless, this delay is short and does not impair evagination resorption, also because all the different effectors involved in PM remodeling can still be recruited by IRSp53. In the case of the I-BAR domain alone, which senses PM curvature and couples it to active Rac1, the delay in recruitment was similar to SH3 mutants but with a stronger impairment in homeostasis recovery. This suggests that the IBAR domain alone, which is expected to be already bound to PM to a certain extent before evagination formation (67), keeps on aggregating at the curved structures in accordance with the sensing mechanism proposed for BAR domains (42, 51), but evagination flattening is impaired by the lack of remaining domains.

Taken together, these data indicate that the recruitment of IRSp53 to the mechanically induced bud-shaped evaginations is necessary for the PM to be successfully remodeled after stretch. The efficiency in the recruitment of this protein ultimately determines the ability of the cell to set in place the fast mechanism mediating PM flattening in response to the physical perturbation.

### Actin polymerization is driven by Rac1 and Arp2/3 activation

Our results point at a role of active Rac1 and further interaction with NPFs to successfully perform PM homeostasis recovery after stretch. Previous work on PM ruffling showed that IRSp53 couples Rac1 to the activation of the WAVE Regulatory Complex (WRC), and the subsequent nucleation of branched actin filaments mediated by Arp2/3 (68–70). However, activation of Arp2/3 downstream of IRSp53 can also be mediated by Cdc42 and N-WASP (41, 56, 59) and, additionally, IRSp53 can coordinate the action of formins mDia1 and mDia2, which drive actin polymerization related to filopodia formation (48, 71). Finally, PM reattachment to the actin cortex may also rely on contractile mechanisms mediated by myosin and not only actin polymerization, as in the case of blebs (72). To discriminate between these mechanisms, we treated IRSp53^-/-R^ cells with different inhibitors and examined evagination resorption after compression. First, cell treatment with 10 μM of the N-WASP inhibitor Wiskostatin (73) reduced filopodia number as expected (74) (Supp. Figs. 4A and B), but did not modify evagination resorption (Figs. 5A, E and I and Supp. Video SV19). Of note, this is consistent with our finding that evagination resorption is not impaired in I268N-CRIB mutant condition in which IRSp53 interaction with Cdc42 is impaired. Second, treatment with 15 μM of the formin inhibitor SMIFH2 (75) reduced the number of filopodia as expected (76) (Suppl. Fig. 4C and D), but did not affect evagination resorption either (Figs. 5B, F and J and Supp. Video SV20). Third, treatment with 10 μM of the myosin II inhibitor Para-nitroblebbistatin (77) affected the integrity of stress fibers as expected (78), (Supp. Fig. 4E) but, again, did not impair evagination resorption (Figs. 5C, G and K and Supp. Video SV21), highlighting that actin polymerization alone is sufficient to drive PM flattening. Consistently and more importantly, treatment with the Arp2/3 inhibitor CK-666 (79) significantly impaired evagination resorption in comparison to DMSO treated controls (Figs. 5D, H and L and Supp. Video 22).

**Fig. 5:**
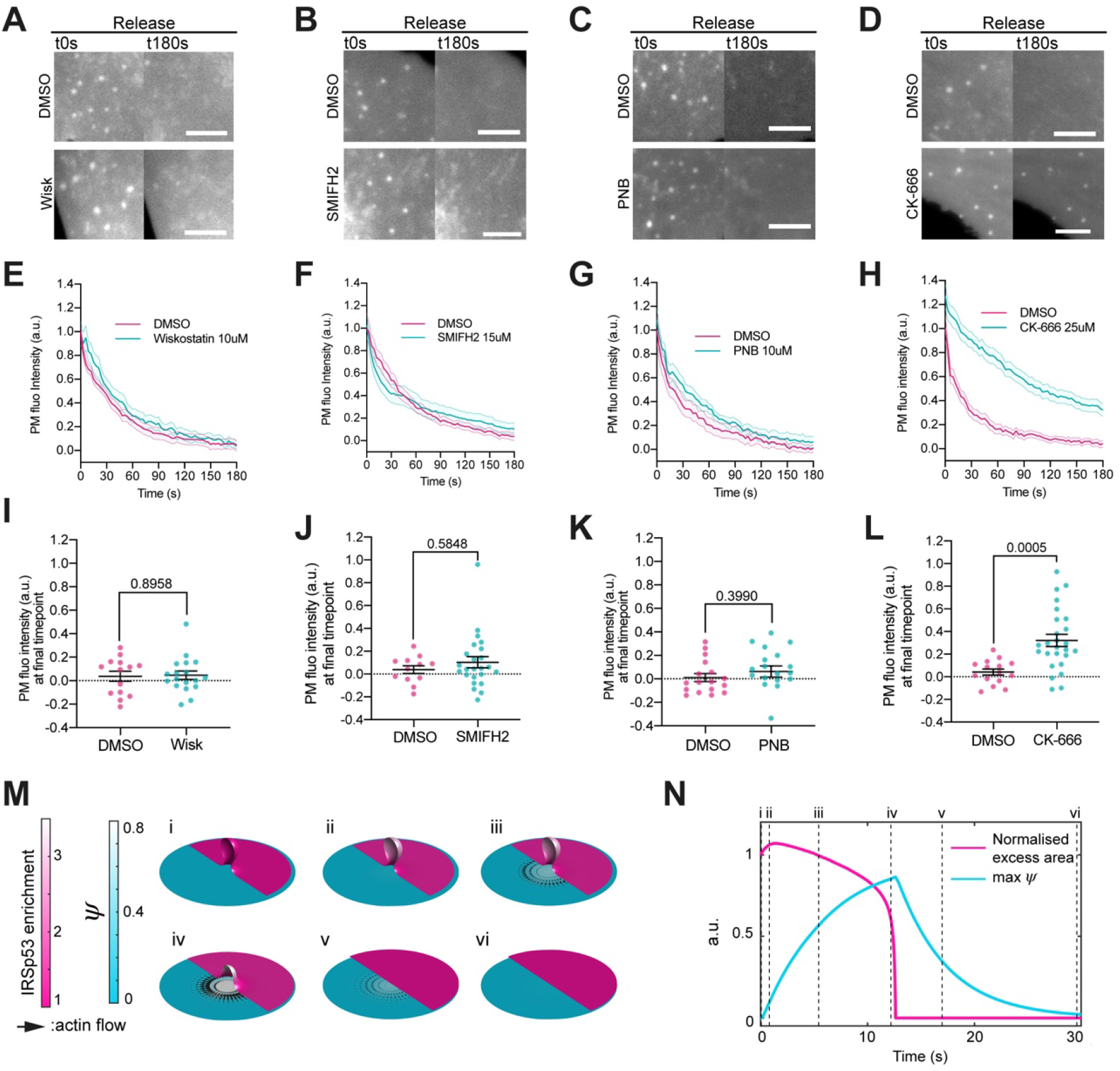
IRSp53 organizes actin polymerization via Arp2/3 activation. **(A-D)** Images after stretch release of PM evaginations, for IRSp53^-/-R^ cells treated with either vehicle (DMSO) or 10 μM Wiskostatin, 15 μM SMIFH2, 10 μM PNB, and 25 μM CK-666, respectively. Scale bars are 5μm. PM is marked with EGFP-membrane. **(E-H)** Corresponding dynamics of PM evaginations. **(I-L)** Differences in PM fluorescence intensity at the final timepoint of acquisition (t180 s after stretch) between DMSO treated control cells and drug treated cells. Statistical significance was assessed through unpaired T-test for CK-666 and PNB against their respective controls, and Mann-Whitney test for SMIFH2 and Wiskostatin against their respective controls. For Wiskostatin, N= 18 and 14 cells, SMIFH2, N = 24 and 12 cells, PNB, N= 19 and 17 cells and CK-666, N= 26 and 15 cells from 3 independent experiments for all cases. **(M)** Dynamics of the model of chemo-mechanical signaling, showing the local enrichment of IRSp53 from a baseline value of 1 (magenta, right side of images) and the concentration of an actin regulator ψ (cyan, left side of images). After the formation of the evagination (i), IRSp53 becomes enriched within 1 second (ii), which triggers the local increase in the concentration of actin regulator ψ over about 10 seconds (iii), thus creating a tension gradient and subsequent centrifugal cortex flow dragging and flattening the membrane (iv, v). Once planarity is restored, the IRSp53 domain rapidly disassembles, the actin regulator recovers its steady-state, and the flow ceases (v, vi). The radius of the membrane patch is 150 nm. (**N**) Corresponding quantifications of PM excess area contained in the evagination (where 0 corresponds to a flat membrane patch) and actin regulator concentration ψ, timepoints corresponding to configurations shown in N are indicated in roman numerals. Both quantifications are normalized to a maximum of 1. Data show mean ± s.e.m.

Thus, evagination resorption upon compression involves the recruitment of IRSp53, leading to actin polymerization in a myosin-independent and Arp2/3-dependent manner. IRSp53 has been described to indirectly promote Arp2/3-mediated actin polymerization acting both as an upstream (80) and downstream regulator of the small GTPase Rac1. To verify whether this was the case in our system, we examined evagination resorption after overexpressing constitutively active (G12V) and dominant negative (T17N) forms of Rac1. Confirming the involvement of Rac1, the expression of Rac1-G12V accelerated evagination resorption significantly whereas Rac1-T17N slowed it down in NHDF (Supp. Figs. 5A, B and C). Finally, and further showing that Rac1 activation is sufficient to trigger evagination resorption, overexpression of constitutively active Rac1-G12V drastically accelerated evaginations resorption even in the background of IRSp53^-/-^ cells (Supp. Fig. 5D, E and F), consistent with an ancillary/modulatory role of IRSp53 in mediating Rac1-dependent activation of the WAVE/Arp2/3 complexes.

### A mechanical mechanism for actin-mediated evagination flattening

Previous work on IRSp53-mediated actin polymerization described the formation of out-of-plane protrusions in the form of filopodia or lamellipodia (*41, 48, 49, 51, 62, 80*). The physical mechanism supporting further protrusion relies on the natural notion that polymerization induces out-of-plane forces on the PM (*81*), which in the case of polymerization by Arp2/3 should push outwards, or at least stabilize protrusions (*82*). At larger scales, polymerization of an actin cortex retracts and flattens cellular blebs, but this mechanism depends on myosin contractility (72), and hence is not applicable here. In contrast, our results show a novel flattening rather than protruding response. To propose a plausible mechanism, we developed a theoretical model coupling the PM and the actin cortex (see methods). We hypothesized that, rather than out-of-plane forces, flattening may be the result of in-plane actin flows around evaginations. We thus approximated the actin cortex as a flat 2D active gel. In this model, the PM is adhered to the underlying cortex from which it can delaminate, and experiences frictional in-plane forces proportional to relative slippage (28). This is coupled to our previous model describing interactions between the PM and curved proteins (*83*). We coarse-grained the signaling pathway triggered by IRSp53 localization and leading to actin polymerization through a regulator species with normalized areal density *ψ*, which is produced beyond a threshold in IRSp53 enrichment, degraded, and transported by diffusion, with dynamics on time-scales comparable to those of actin dynamics. The effect of this regulator is to locally favor actin polymerization by the Arp2/3 complex, and hence bias the competition between a formin-polymerized contractile network component and a branched extensile component (*84, 85*). We thus modelled the mechanical effect of local polymerization by locally reducing contractility.

Our model predicted that curvature-sensitive IRSp53 molecules became enriched in the evagination within a second after its formation. This led to recruitment of the regulator species *ψ*, resulting in a tension gradient in the vicinity of the evagination. In turn, this induced a centrifugal cortical flow, which frictionally dragged the membrane outwards until flattening. In the absence of curvature, the IRSp53-enriched domain dissolved, the regulator species recovered its uniform baseline, and the cortex recovered its quiescent steady-state (Fig. 5M and N). Whereas predicted actin flows occur at a scale well below the diffraction limit and can therefore not be observed experimentally, the predicted relative trends of PM and regulator densities qualitatively match our experimental observations when comparing PM and actin (Fig. 2B) or ezrin (Fig. 2E). We note that in the real system, the proposed mechanism based on in-plane actin flows and cortex-PM friction should compete with the classical mechanism based on out-of-plane forces. This may explain why resorption dynamics in experiments (Fig. 2B and E) were significantly longer and less abrupt than those predicted by the model (Fig. 5N). Predictions are also consistent with our observation that evagination resorption is impaired when inhibiting Arp2/3 (Fig. 5D) but not myosin or formin activity (Fig. 5B and C). Indeed, the mechanism is based on a local gradient in extensile versus contractile behavior around the evagination, so it should depend on Arp2/3 (which acts locally at the evagination) and not on formin or myosin, which would regulate overall contractility levels and not specifically local gradients. Thus, our model suggests a chemo-mechanical signaling system that autonomously restores homeostasis of membrane shape.

## Discussion

Our work shows that stretch-compression cycles generate evaginations on the apical PM of the cells with a size on the 100 nm scale, compatible with the sensing range of IBAR proteins (47, 51). Further, we demonstrate the recognition of this curved templates by the curvature-sensing protein IRSp53. The role of IRSp53 is not due to general cell-scale effects, such as lamellipodial extension (44, 57) or endocytosis. Indeed, cell spreading after the stretch-compression cycle was not affected by IRSp53 (Supp. Fig. 2). Regarding endocytosis, IRSp53 has been described to regulate the CLIC-GEEC endocytic pathway (50), which is in turn activated upon cell compression (18). However, the I268N-CRIB and 4KE-IBAR IRSp53 mutants strongly impaired endocytosis (50), but fully rescued evagination resorption (Fig. 3F), showing that IRSp53 affects both phenomena through different mechanisms. Further supporting this possibility, the resorption of low-curvature VLDs formed upon transient exposure of cells to hypo-osmotic media, a treatment which also activates CLIC-GEEC endocytosis (18), was not affected by IRSp53.

Thus, our findings demonstrate a novel mechanosensing mechanism: upon cell compression, cells are known to use caveolae formation (22) and the CLIC-GEEC endocytic pathway (18) to store material from the PM and recover resting tension. On top of this, we demonstrate a new event at the local scale, which restores PM shape perturbations induced by mechanical stimulation. This event involves the progressive flattening of the PM and not its scission, which would have involved an abrupt loss of evagination fluorescence (and the appearance of fluorescent membrane vesicles) which we never observed in experiments. To achieve such flattening, cells employ the IRSp53-Rac1-Arp2/3 network, well described to polymerize actin in the context of lamellipodia extension or ruffling (*86, 87*), and revisited here to describe its action in response to physical perturbations. In this regard, we describe a novel mechanism, and biophysical framework, in which Arp2/3 mediated actin polymerization can lead to membrane flattening rather than protrusion.

While stretch is often studied separately from subsequent compression provoked by its release (*88-90*), here we put in relevance the coupling between the two at the single cell level. Such coupling takes place for instance in heart beating, breathing, the musculoskeletal system, or in many developmental scenarios. Thus, and although this remains to be explored, our mechanism could be relevant in events such as the fast compressions of cells embedded in connective tissues (*91*), or apical expansion and contractions of amnioserosa cells during dorsal closure in Drosophila embryos (*92*), among many others. In conclusion, our findings reveal a new mechanosensing mechanism explaining how PM detects physical stimuli at a local, sub-μm scale, and further coordinates a response allowing for quick adaptation to a changing environment.

## Materials and Methods

### Cell culture, expression vectors and reagents

NHDF were purchased from Lonza (CC-2511) and cultured in DMEM without pyruvate (ThermoFisher 41965-039) supplemented with 10% FBS (v/v), 1% penicillin-streptomycin (v/v) and 1% Insulin-Transferrin-Selenium (v/v) (ThermoFisher 41400045). IRSp53^-/-^ MEF infected with an empty pBABE or a pBABE-IRSp53-WT retroviral vector were generated by G. Scita (IFOM, Milan) as previously described (52-54), leading to a cell line that we noteIRSp53^-/-R^. The culture was maintained in DMEM supplemented with 1 % penicillin-streptomycin (v/v) and 1 μ/mL puromycin to selectively maintain cells expressing the selection vector. CO2 independent media (ThermoFisher 18045088) was used for microscopy imaging and was supplemented with 10μg/mL of rutin (Sigma R5143) to prevent photobleaching (*93*). mCherry, EGFP and EYFP membrane markers contained a fusion protein consisting in one of the three fluorophores coupled to the 20 last amino acids of Neuromodulin which is post-translationally palmitoylated and targets the fluorophore to PM (20). IRSp53 60950 shRNA and control Non-Targeting shRNA were purchased from Sigma Mission for viral transfection and stable cell line creation. mEmerald-Ezrin was from Addgene (#54090). EGFP-IRSp53-FL (62), EGFP-IRSp53-4KE, EGFP-IRSp53-I268N (50) and EGFP-IRSp53-I403P (52) contained isoform 2 of the murine protein either wild type or carrying the mentioned mutations in the pC1-EGFP backbone. EGFP-IRSp53-W413G,EGFP-IRSp53-ΔIBAR and EGFP-IBAR (52) where created based on the sequence of isoform 4 of the human protein inserted in the pC1-EGFP backbone. A point mutation was included in the SH3, the first 312 amino acids were removed in the case of the ΔIBAR and the first 250 amino acids were expressed to obtain the IBAR-domain. The dominant constitutively active Rac1-G12V and the dominant negative Rac1-T17N were described previously (*94*). Actin was marked using the mammalian expression vector encoding the cytoskeleton marker Actin-VHH fused to either or RFP or GFP2 and commercially sold as Actin-Chromobody^®^ (Chromotek)

On the day prior to the experiment, cells were transfected by electroporation with the selected plasmids using the NeonTM Transfection System (Invitrogene) following the protocol provided by the company. CK-666 was purchased from Merck (Ref 182515), SMIFH2 was from Abcam (ab218296), Wiskostatin was bought from Sigma (W2270) and Para-Nitro-Blebbistatin was from Optopharma (DR-N-111). All compounds were diluted in DMSO and conserved according to manufacturer’s instructions. On the day of the experiment, drugs were diluted in culture media, filtered through a 0,22 μm filter and warmed up to 37°C prior to addition to the culture. Cells were treated with 25 μM of CK-666 for 30 min, 10 μM of PNB for 30-40 min and 10 μM Wiskostatin or 15 μM SMIFH for 1 h prior to the experiment.

### PDMS membrane fabrication

The stretchable PDMS membranes were prepared as described in (20). To produce a patterned support to further obtain patterned-PDMS membranes PMMA dishes were plasma cleaned for 20 min and warmed up to 95°C for 5 min. After cooling down using a nitrogen gun, SU 2010 resin was spinned on top of the dish to create a 10 μm layer and prebaked 2,5 min at 95°C. Dishes were then placed on a mask aligner and exposed for 7,5 s in presence of the designed acetate mask. After post-baking for 3,5 min at 95°C, the pattern was revealed for 1 min and subsequently extensively washed with isopropanol and verified under the microscope. Finally, PMMA dishes were silanized by 30 s plasma cleaning activation followed by 1 h silane treatment under vacuum. Standard or patterned membranes were mounted on metal rings of our customized stretch system, cleaned, sterilized, and coated with 10 μg/ml fibronectin (Sigma) overnight at 4°C prior to experiments. Patterns were designed as a grid with letters and numbers to allow for correct orientation.

### Stretch and osmolarity experiments

After overnight fibronectin coating, PDMS membranes were quickly washed and 3000 cells were seeded on top and allowed to spread for 45min to 1h in the incubator. Then, rings were mounted on the stretch device coupled to the microscope stage, vacuum was applied for 3 min to stretch the membrane, and then vacuum was released to come back to the initial shape as described in (20). Calibration of the system was done to adjust the vacuum applied to obtain 5 % stretch of the PDMS surface. Hypo-osmotic shocks were performed by exposing cells during 3 min to CO2 independent medium mixed at 50% with de-ionized water in which the concentrations of Ca^+2^ and Mg^+2^ had been corrected. Iso-osmotic medium was added after the 3 min incubation period.

### Scanning electron microscopy experiments

Cells were prepared as explained in the previous section. Right after stretch release, the sample was fixed in 2.5 % glutaraldehyde EM grade (Electron Microscopy Sciences 16220) plus 2 % PFA (Electron Microscopy Sciences 15710-S) diluted in 0.1 M PB buffer at 37°C for 1 h. Samples were then washed 4x for 10 min in 0.1 M Phosphate Buffer (PB) and imaged with epifluorescence microscopy as described below to acquire fluorescence images of the cell PM. PDMS membranes were then cut into 1×0.5 cm rectangles in which the pattern was centered and placed on top of 12 mm coverslips for further processing. Dehydration was carried out by soaking samples in increasing ethanol concentrations (50, 70, 90, 96 and 100 %). After this, samples were critical point dried and covered with a thin layer of gold to be imaged.

### Transmission electron microscopy experiments

Cells were fixed, washed and PDMS membranes were cut and mounted as for SEM imaging. After this, samples were postfixed with 1% OsO4 and 0.8 % K_3_Fe(CN)_6_ for 1 h at 4°C in the dark. Next, dehydration in increasing ethanol concentrations (50, 70, 90, 96 and 100%) was done. Samples were then embedded in increasing concentrations of Pelco^®^ EPONATE 12TM resin (Pelco 18010) mixed with acetone. 1:3 infiltration was done for 1 h then 2:2 for 1h and finally 3:1 overnight. On the next day, embedding was continued with EPON12 without catalyzer for 3×2 h washes and then overnight. Last, samples were embedded in EPON12 plus catalyzer DMP-30 (Pelco 18010) for 2×3 h. To finish, blocks were mounted and polymerized for 48 h at 60°C. PDMS membrane was next peeled off and ultrathin sections were cut and mounted on grids for imaging.

### APEX labelling for TEM imaging

Two days prior to the experiment, cells were co-transfected by electroporation with mKate2-P2A-APEX2-csGBP (Addgene #108875) and EGFP-IRSp53-FL in a 3:1 ratio, using the Neon™ Transfection System (Invitrogene) following the protocol provided by the company. Before seeding, cells were sorted for double positive mKate and GFP fluorescence, excluding very high and very low transfection levels. Cells were subsequently seeded and stretched in the same conditions as explained in the stretch experiments section. Right after stretch release, the sample was fixed in 2.5 % glutaraldehyde EM grade (Electron Microscopy Sciences 16220) diluted in 0.1 M Cacodylate buffer at 37°C for 10 min, followed by incubation on ice for 50 min in presence of the fixative. All subsequent steps were performed on ice. The sample was washed 3 times with cold 0.1 M Cacodylate buffer, and next cut into 1×0.5 cm rectangles containing the fixed cells. Cells were washed for 2 min with a fresh cold 1 mg/ml 3,3’-diaminobenzidine (DAB) (tablets, Sigmafast, D4293) solution in 0.1 M Cacodylate buffer. Cells were immediately incubated with a fresh cold 1 mg/ml DAB solution in cold 0.1 M Cacodylate buffer supplemented with 5,88 mM hydrogen peroxidase (PERDROGEN™ 30% H2O2, 31642, Sigma). The samples were washed 3 times with cold 0.1 M Cacodylate buffer, and subsequently incubated for 30 minutes with cold 1% OsO4. Dehydration, resin embedding, and block mounting was done as described in the TEM experiments section.

### Image acquisition

Fluorescence images were acquired with Metamorph software using an upright microscope (Nikon eclipse Ni-U) with a 60x water dipping objective (NIR Apo 60X/WD 2.8, Nikon) and an Orca Flash 4.0 camera (Hamamatsu). Fluorophore emission was collected every 3s. Cells were imaged in a relaxed state and then for 3 min at 5% stretch, and for 3 min during the release of stretch. SEM images were taken using the xTm Microscope Control software in a NOVA NanoSEM 230 microscope (FEI Company) under the high vacuum mode using ET and TL detectors to acquire high and ultra-high resolution images of the cell surface. TEM Samples were observed in a Jeol 1010 microscope (Gatan, Japan) equipped with a tungsten cathode in the CCiTUB EM and Cryomicroscopy Units. Images were acquired at 80 kv with a CCD Megaview 1kx1k.

### Fluorescence analysis

All images used for time course analysis were aligned using the Template Matching plugin from Fiji to correct the drift. To assess the evolution of PM evaginations, VLDs or the different marked proteins, their fluorescence was quantified. To ensure that we only considered the fluorescence of structures induced by stretch or osmotic shocks, the analysis was carried out in regions devoid of visible endomembrane structures before the application of stretch or osmotic shocks. For each evagination, we calculated the integrated fluorescence signal of a small region of interest containing the evagination (*I_evag_*), the integrated fluorescence signal of a neighboring region of interest of the same size and devoid of any structures (*I_PM_*), the integrated fluorescence signal of the entire cell (*I_cell_*) and the integrated fluorescence signal of a background region of the same size as the cell (*I_BG_*). Then, the final evagination signal *I_final_* was computed as:

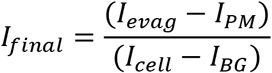

The numerator of this expression corrects evagination fluorescence so that only the signal coming from the evagination itself and not neighboring PM is quantified. The denominator normalizes by total cell fluorescence, and also accounts for progressive photobleaching. All control curves were normalized to 1 (maximal fluorescence after stretch release) and the rest of the data represented in the same graph were normalized to the control. Exceptionally, actin and ezrin curves were normalized to 0.5 (maximal fluorescence after the release of stretch) for visualization purposes. To quantify the degree of resorption of the evaginations, as the experimental data could not always be fitted with single exponential decay curve, we adopted the strategy of comparing the residual fluorescence intensity of the PM marker at the las timepoint of acquisition (t180s), on which statistical analysis can be performed. Full reabsorption of evaginations leads to a complete return to fluorescent baseline (≈0), while presence of a residual fluorescence indicates non-reabsorbed evaginations. Lag time was calculated by identifying the maximum intensity timepoints in the protein and PM channels, and subtracting them to obtain the time between the two events.

### Area analysis

To compute the changes in cell area with time after stretch, automated area analysis for each timepoint was done using CellProfiler (*95*) (https://cellprofiler.org/). To calculate the time constant (k) of each experimental curve, data was fitted to a one-phase decay (for time course dynamics of PM evaginations, VLDs and protein markers) or one-phase association equation (area analysis after stretch) using GraphPad and k was extracted from the fittings to be further compared by statistical analysis.

### Quantification of number and PM Area % stored by evaginations

3 regions of different parts of the cell where randomly chosen from every cell at the timepoint t0s (right after the release of stretch) and the number of evaginations was manually counted by comparing the analyzed images with the images of the cell during stretch, to discard PM structures not formed by stretch-release. For stored area calculation, the membrane area fraction mf contained in evaginations was estimated as:

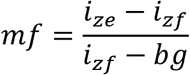

Where i_ze_ is the average fluorescence intensity of a cell zone (containing evaginations), i_zf_ is the average fluorescence intensity of a neighbouring flat patch of membrane (small enough so that it does not contain any evaginations), and bg is the average intensity of background. For each cell, this was done for 3 random regions containing evaginations.

### Fluorescence and SEM correlation

Images of the fixed sample were acquired in fluorescence and brightfield and positions of the imaged cells in the pattern were noted down. Sample was then processed for SEM imaging and the same cells were found by manually following their location on the pattern and visual verification was done to check for correct matching. Fluorescent and SEM images were then aligned by using the BigWrap plugin on Fiji.

### Statistical analysis

In the case of data following a normal distribution, T-test or ANOVA was done depending on whether there were 2 or more datasets to compare. For data not following normal distributions, Mann-Whitney or Kruskal-Wallis test were applied depending on whether there were 2 or more datasets to test. All data are shown as mean ± SEM. Specific P and N values can be found in each one of the graphs shown in the figures.

### Theoretical Model

To understand the physical mechanism leading to the active flattening of membrane evaginations caused by compression of the PM, we focused on a single evagination and described it mathematically under the assumption of axisymmetry. We modelled the membrane as locally inextensible thin sheet with bending rigidity *k* =20 *k_B_T* using the Helfrich model and accounted for the viscous stresses due to membrane shearing with membrane 2D viscosity *η_m_* = 3 · 10^-3^pN s/μm (*28, 30, 96*). We modelled the cortex as a 2D planar active gel adjacent to the membrane. We thus ignored the out-of-plane protrusive forces caused by localized actin polymerization at evaginations enriched in IRSp53, which in a classical view can lead to further protrusion rather than flattening (*55*). Instead, we focused on the in-plane effect of localized actin polymerization to explain active flattening. In the actual system, we expect both effects to compete.

To model the interaction between the membrane and the cortex, we considered an adhesion potential depending on the distance between the membrane and the cortex enabling decohesion with an adhesion tension of *γ* = 1.5 · 10^-5^ N/m (*30*), (Supp. Fig. 6). We also considered in-plane frictional tractions between the membrane and the cortex proportional to their relative velocity, *τ* = *μ*(*v_m_* – *v_c_*) where *v_m_* is the membrane velocity, *v_c_* is the cortex velocity, and *μ* is a friction coefficient, which we took as *μ* = 20 nN s/μm^3^ (*28*).

We generated evaginations with dimensions comparable to those in (Fig. 1) by laterally compressing an adhered membrane patch of radius *R*_0_ as discussed in (*30*). We considered *R*_0_ = 150 nm, consistent with the typical separation between evaginations (Fig. 1C). After formation of the evagination, we applied at the boundary of our computational domain the surface tension required to stabilize the evagination, consistent with the long-time stability of such compression-generated evaginations of the PM when cellular activity is abrogated (*20*).

We then considered the model in (*83*) to capture the interaction between an ensemble of curved proteins (IRSp53) and a membrane. In this model, proteins are described by their area fraction *ϕ*. We fixed the chemical potential of such proteins at the boundary of our computational domain, corresponding to a relatively low area fraction of proteins, 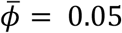. We set the saturation coverage to *ϕ*^max^ = 0.35 due to crowding by other species but in our calculations, coverage did not come close to this limit. We considered an effective surface area per dimer of 300 nm^2^. In this model, the curvature energy density of the membrane-protein system is given by 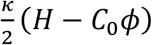 where H is the mean curvature and *C*_0_ is a parameter combining the intrinsic curvature of proteins and their stiffness (*83*). We took *C*_0_ = 3 · 10^-3^ nm^-1^, which lead to curvature sensing but no significant protein-induced membrane reshaping. With a protein diffusivity of 0.1 μm^2^/s, we obtained protein enrichments on the evagination of about 3-fold within 0.5 s.

To model in a coarse grained manner the signalling pathway triggered by IRSp53 localization and leading to actin polymerization, we considered a regulator species given by a normalized surface density *ψ*, which was produced with a rate depending on IRSp53 enrichment and given by 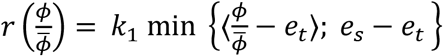, where *e_t_* is a threshold IRSp53 enrichment for signaling, *e_s_* is an enrichment saturation threshold beyond which the production of *Ψ* saturates, and 〈*a*〉 is 0 if *a* < 0 and a otherwise. We considered *e_t_* = 2, *e_s_* =3 and *k*_1_ =1 s^-1^. This regulator was degraded with rate *K*_2_*ψ*, with *k*_2_ = 1 s^-1^ and diffused with an effective diffusivity of *D* = 0.1 o 10^-3^ μm^2^/s, much smaller than that of membrane proteins since the regulator is viewed as an actin-binding species. In polar coordinates, the governing equation for the transport of this regulator is thus

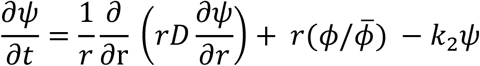

This equation results in a region enriched with *ψ*, co-localizing with the evagination, and reaching a maximum value of about 1 within about 10 s, comparable to the typical times of actin dynamics. Not being a detailed description of a specific network, the details of this model for *ψ* are not essential. The key points are that the production of *ψ* is triggered by IRSp53 enrichment, and that *k*_1_, *k*_2_ and *D* are such that over the time-scales of actin dynamics (significantly slower than those of IRSp53 enrichment) a region of high *ψ* develops close to the evagination.

The effect of this regulator is to locally favour actin polymerization by the Arp2/3 complex. The cortex can be viewed as a composite system of interpenetrating actin networks, one polymerized by formins leading to linear filaments and producing contractile forces through the action of myosins and other crosslinkers, and one polymerized by the Arp2/3 complex, with a branched architecture and producing extensile forces by polymerization (*84*). Combining these two effects, the net active force generation in the actin cortex is contractile. These two networks compete for actin monomers (*85*), and hence a local enrichment in the regulator leading to enhanced polymerization of the branched network should bias this competition and locally lower contractility in the vicinity of the evagination. In turn, the resulting contractility gradient should generate an in-plane centrifugal cortical flow, which if large enough, might drag the membrane outwards due to frictional forces and actively flatten the evagination.

To model such actin flow, we considered simple active gel model where the cortical velocity *υ*_c_ is obtained by force balance between viscous and active forces in the cortex, and given by

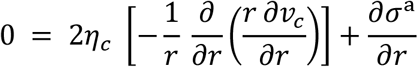

where η_c_ is the viscosity of the cortex and *σ^a^*(*ψ*) is the active tension, which we assume to be a function of the regulator ψ. We note that we neglect in the equation above the force caused by friction between the membrane and the cortex as they slip past each other. This is justified because the hydrodynamic length for the cortex is in the order of microns and above, and hence in the smaller length-scales considered here viscosity dominates over friction. In our calculations, we took 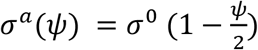, so that active tension is approximately halved near the evagination when the normalized regulator density *ψ* reaches about 1 and is equal to *σ*^0^ far away from it. As boundary conditions, we considered *v_c_* (0) = 0 consistent with polar symmetry and 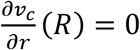, so that at *r* = *R* the stress at the gel is *σ*^0^. We chose *σ*^0^/*η_c_* so that the resulting cortical velocities due to gradients in active tension gradients were of about 0.1 μm/s, comparable to the typical actin velocities due to polymerization in the lamellipodium (*97*).

The formation of the evagination triggered in this model a sequence of chemo-mechanical signaling event restoring autonomously homeostasis of membrane shape and of all the signaling network. Indeed, within a few seconds, IRSp53 became enriched in the evagination by curvature sensing. Then, over a about 10 seconds, the actin regulator *ψ* progressively built up in the vicinity of the evagination, creating a gradient in active tension *σ*, which in turn created a centrifugal cortical flow. This flow frictionally dragged the membrane outward ironing out the evagination. In the absence of curvature, the IRSp53 domain rapidly dissolved and according to Eq. (1) *ψ* dropped to zero everywhere, eventually stopping the cortical flow and thus recovering a homeostatic state with a planar membrane and a quiescent cortex.

We note that our model is consistent with the fact that myosin inhibition does not affect the resorption process. Indeed, myosin inhibition should lower the baseline active tension, *σ*_0_, but should not change the fact that localized polymerization would locally induce and extensile stress, and hence establish a tension gradient and an actin flow.

One important difference between our model and the experiments is that, in our calculations, the evaginations rapidly flattened once the contact angle of the evagination became smaller than 90 degrees, whereas in the experiments, the decay of membrane fluorescence was more gradual over a timescale of 3 minutes. We hypothesize that this may be due to the fact that localized actin polymerization may fill the evagination with branched actin network, which should apply an out-of-plane force competing with the flattening force causing the centrifugal flow and whose material needs to be cleared out even when localized polymerization has stopped. Both of these effects should slow down the resorption process.

## Acknowledgments

We thank V. González-Tarragó for assistance with the stretch system and statistics analysis, L. Rosetti for support with Fiji scripts, I. Granero for helping with CellProfiler pipelines, N. Castro, S. Usieto and A. Menéndez for technical assistance and the members of the P.R.-C. and X.T. laboratories for technical assistance and discussions. We also would like to acknowledge the support given by the Unitat de Criomicroscòpia Electrònica TEM/SEM (Centres Científics i Tecnològics de la Universitat de Barcelona, CCiTUB) and the MicroFabSpace and Microscopy Characterization Facility, Unit 7 of ICTS “NANBIOSIS” from CIBER-BBN at IBEC.

## Funding

Spanish Ministry of Science and Innovation (BFU2015-66785-P to F.T:, PGC2018-099645-B-I00 to X.T., PID2019-110949GB-I00 to M.A., BFU2016-79916-P and PID2019-110298GB-I00 to P. R.-C, and BFU2016-79916-P to XQ) European Commission (H2020-FETPROACT-01-2016-731957) European Research Council (CoG-616480 to X.T. and CoG-681434 to M.A.) Generalitat de Catalunya (2017-SGR-1602 to X.T. and P.R.-C., 2017-SGR-1278 to M.A.)

The prize “ICREA Academia” for excellence in research to M.A. and P.R.-C.

Fundació la Marató de TV3

Obra Social “La Caixa”

IBEC and CIMNE are recipients of a Severo Ochoa Award of Excellence from the MINECO

AC was supported by a FPU fellowship from Ministerio de Educación, Cultura y Deporte (Spain).

Associazione Italiana per la Ricerca sul Cancro AIRC-IG 18621 and 5XMille22759 to GS

The Italian Ministry of University and Scientific Research (PRIN 2017-Prot. 2017HWTP2K to GS)

## Author contributions

Conceptualization: PRC, ALLR, XQ and MA

Methodology: PRC, ALLR, XQ, MIG, GS, AD, FT, XT, RGP and MA.

Investigation: XQ, NW, AC and AM

Visualization: XQ

Supervision: PRC and ALLR

Writing—original draft: XQ

Writing—review & editing: MA, ALLR and PRC.

## Data and materials availability

All data are available in the main text or the supplementary materials.

## SUPLEMENTARY MATERIALS

**Supp. Fig. 1:**
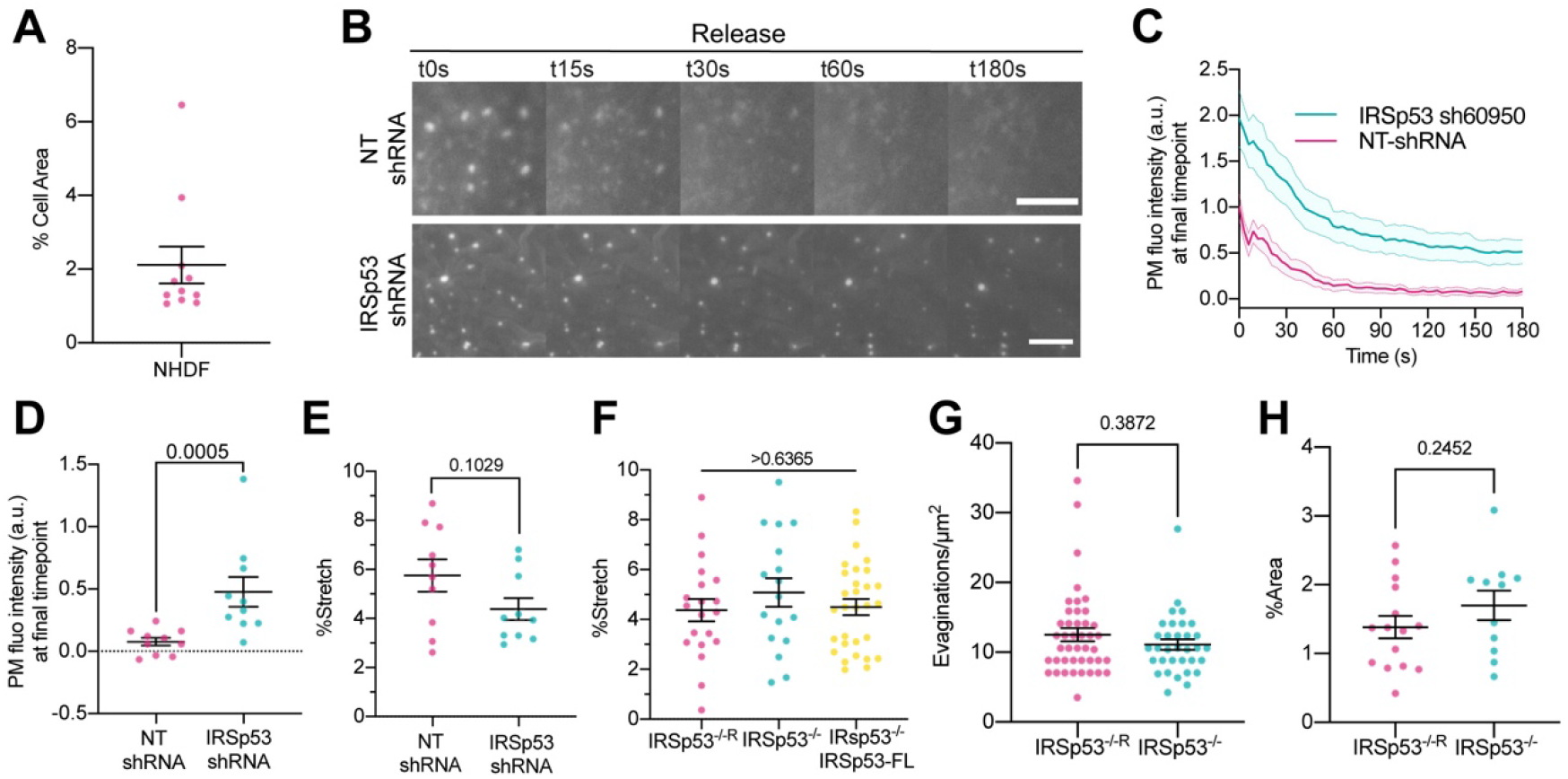
IRSp53 silencing impairs compression-generated PM evagination resorption in NHDF. **(A)** % of cell area stored in PM evaginations after stretch in NHDF. N = 11 from 3 independent experiments. **(B)** Time course images after stretch release of stable NHDF cell lines expressing either a non-targeting (NT) shRNA or an shRNA specifically targeting IRSp53. PM is marked with EGFP-membrane. Scale bars are 5 μm. **(C)** Quantification dynamics of EGFP-membrane tagged PM evaginations after stretch release in NT-shRNA and IRSp53 shRNA expressing cells. **(D)** Differences in EGFP-membrane fluorescence intensity at the final timepoint of acquisition after stretch in the conditions mentioned above. Significance was calculated through Mann-Whitney test. **(E)** Areal stretch experienced by NT-shRNA and IRSp53 shRNA expressing cells under exposure to 7% PDMS membrane nominal stretch. Statistical differences were tested through unpaired T-test. N=8 and 10 cells from 2 independent experiments. **(F)** Areal stretch experienced by IRSp53^-/-R^, IRSp53^-/-^ and IRSp53^-/-^EGFP-IRSp53-FL under exposure to 5% PDMS membrane nominal stretch. Statistical differences were tested through one-way ANOVA. N= 20, 19 and 28 cells from 3, 3 and 5 independent experiments. **(G)** Number of PM evaginations per μm^2^ formed after stretch in IRSp53^-/-R^ and IRSp53^-/-^ MEF. N= 43 and 33 regions from 15 and 11 cells from 3 independent experiments. **(H)** % of cell area stored in PM evaginations after stretch in IRSp53^-/-R^ and IRSp53^-/-^ MEF. N = 15 and 11 cells from 3 independent experiments. Statistical differences were tested through Mann-Whitney test. Data show mean ± s.e.m.

**Supp. Fig. 2:**
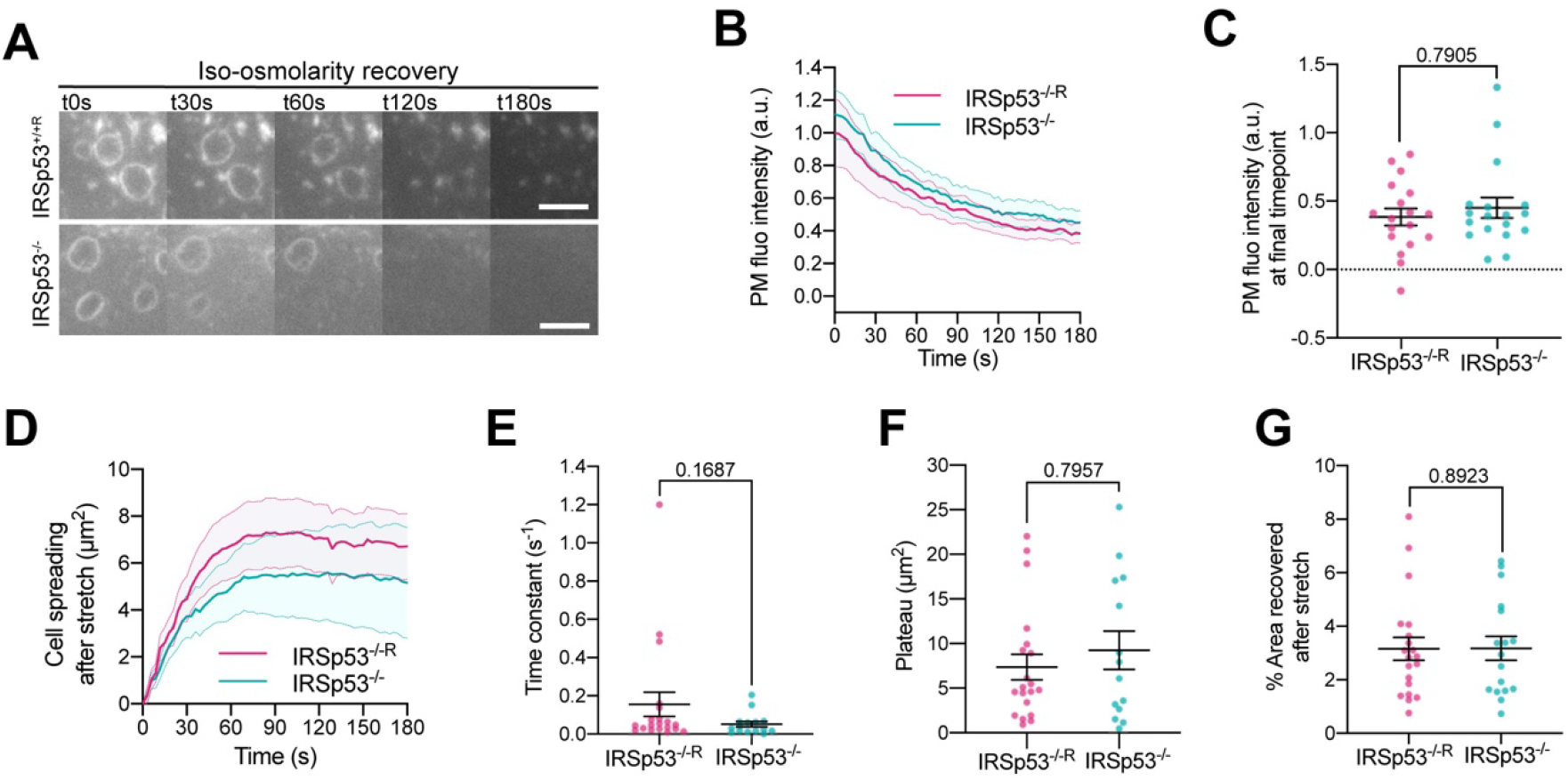
The role of IRSp53 is local and specific to PM evaginations. **(A)** Time course images of VLDs (observed with a pYFP-membrane fluorescent marker transfection) formed by exposing cells to iso-osmolar medium after a transient exposure to a 50% hypo-osmotic medium. Results for IRSp53^-/-R^ and IRSp53^-/-^ cells are shown. Scale bars are 5μm. **(B)** VLDs fluorescence quantification as a function of time. **(C)** Comparison of PM fluorescence intensity of VLDs at the last frame of acquisition (180s after the iso-osmotic medium recovery). Significance was assessed through Mann-Whitney test. N=18 cells from 3 independent experiments. **(D)** Cell spreading during PM recovery phase. 0 = cell area after the release of stretch. **(E)** Comparison of time constants resulting from the exponential fitting of the curves obtained from cell spreading during the recovery phase after stretch. **(F)** Comparison of plateau values resulting from the same exponential fitting. **(G)** Quantification of % of area recovered after stretch. N=20 and 17 cells from 4 independent experiments. Statistical significance was assessed through Mann-Whitney test. Data show mean ± s.e.m.

**Suppl. Fig. 3:**
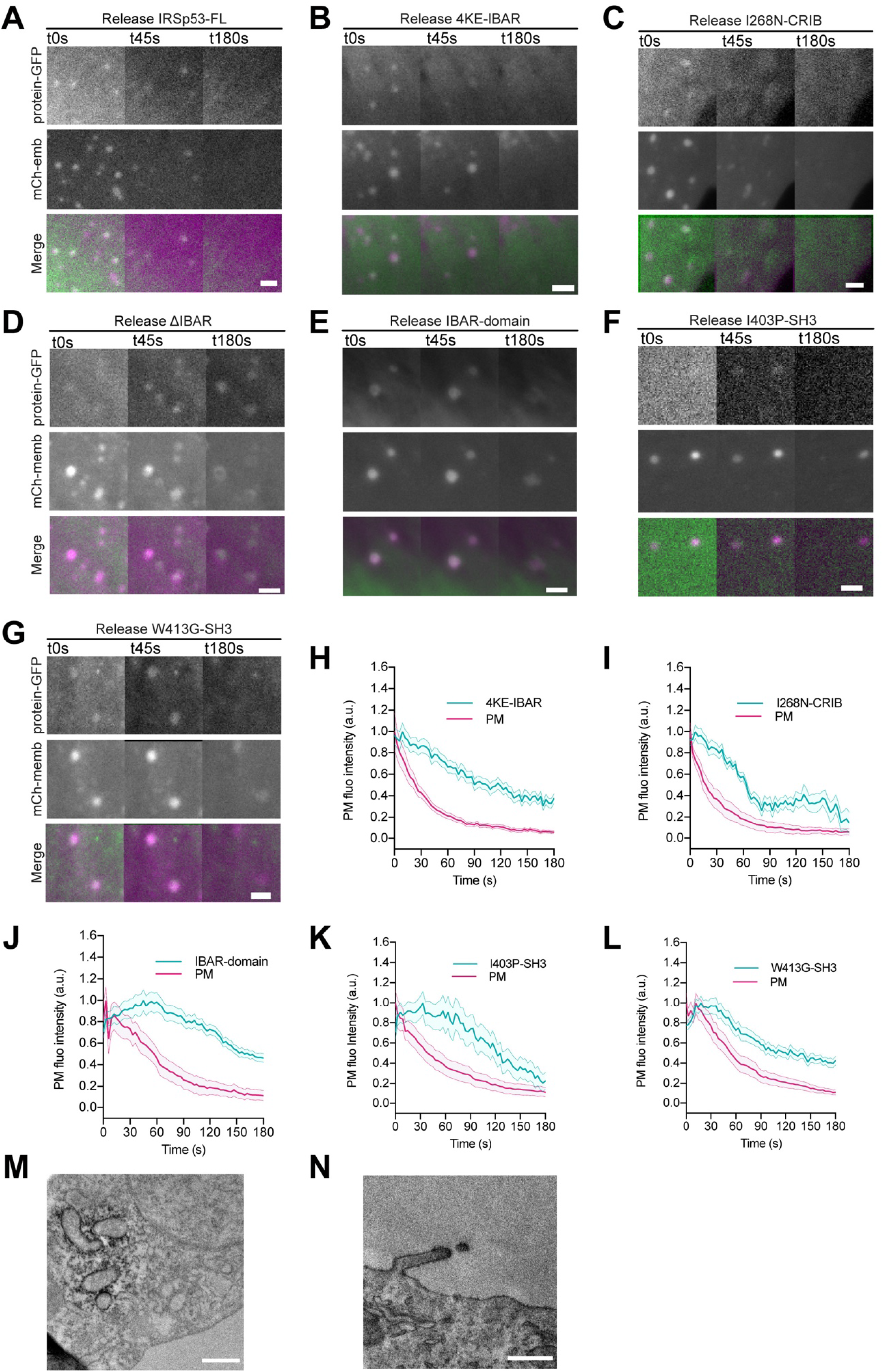
Additional data on IRSp53 mutants. **(A-G)** Images of IRSp53^-/-^ cells after stretch release transfected with mCh-membrane and either the FL form of IRSp53 or different mutant forms of the protein coupled to EGFP. Scale bars are 2μm. **(H-L)** Corresponding dynamics of PM evaginations upon stretch release quantified through mCh-membrane or GFP coupled to the different IRSp53 mutants. N= 15, 13, 14, 9 12 and 16 cells from 3, 3, 2, 3, 3 and 3 independent experiments. **(M-N)** TEM images of IRSp53^-/-^ cells co-transfected with csAPEX2-GBP together with **(M)** mito-GFP or **(N)** EGFP-IRSp53-FL. APEX staining can be observed around mitochondria (M), in the tips of filopodia and up to some extent in the PM of EGFP-IRSp53-FL transfected cells (N), as expected. Scale bars are 500 nm. Data show mean ± s.e.m.

**Suppl. Fig. 4:**
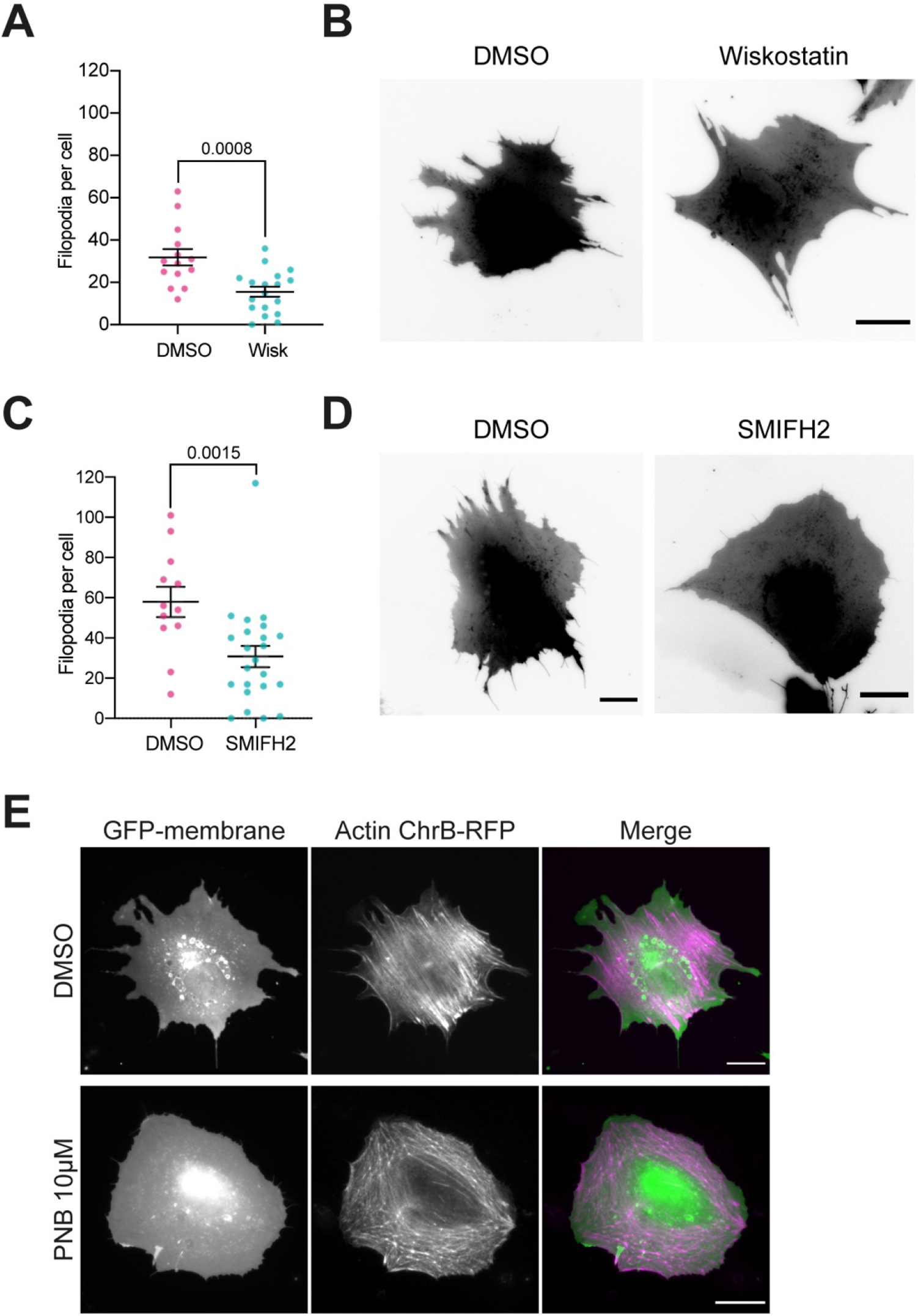
Controls of drug treatment in IRSp53^-/-^R MEF. **(A)** Number of filopodia per cell in 10 μM Wiskostatin or vehicle (DMSO) treated cells. Compound was incubated for 30 min at 37°C before experiments. N= 18 and 14 cells respectively from 3 independent experiments. Statistical significance was assessed through unpaired T-test. **(B)** Corresponding images of GPF-membrane transfected cells. **(C)** Number of filopodia per cell in 15 μM SMIFH2 or vehicle (DMSO) treated cells. Compound was incubated for 1 h at 37°C before experiments. N=24 and 13 cells form 4 independent experiments. Statistical significance was assessed through Mann-Whitney test. **(D)** Corresponding images of GPF-membrane transfected cells. **(E)** IRSp53^-/-R^ MEF after 30 min incubation at 37°C with either 10 μM PNB or vehicle (DMSO). Cells were transfected with GFP-membrane and Actin Chromobody-RFP to mark both PM and actin. For all images scale bar is 20 μm. Data show mean ± s.e.m.

**Suppl. Fig. 5:**
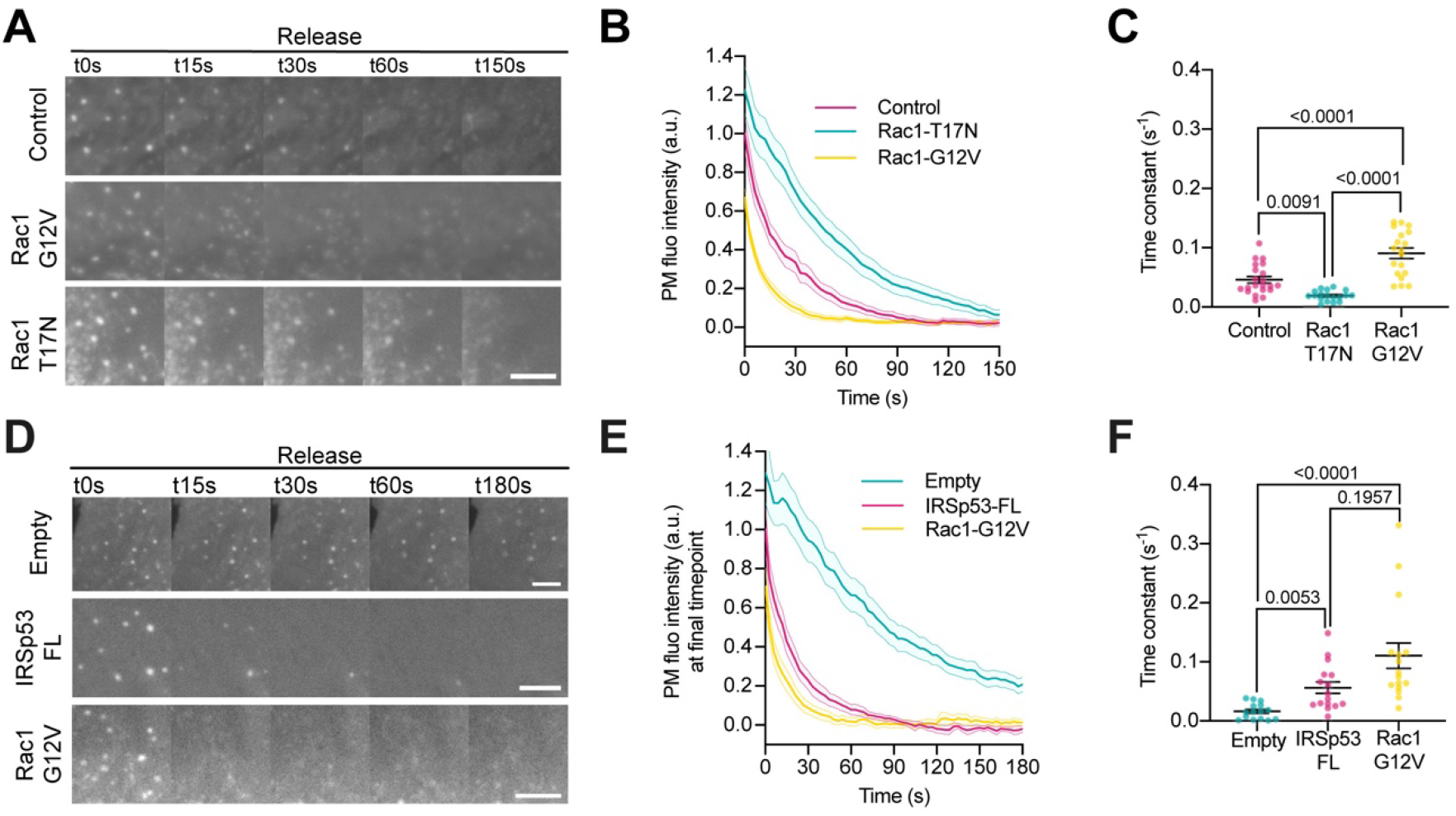
Rac1 is involved in PM remodeling upon stretch. **(A)** Time course images of PM evaginations after stretch release on NHDF expressing a PM marker alone, or a PM marker plus either a constitutively active (G12V) or a dominant negative (T17N) form of Rac1. PM was tagged with GFP-membrane marker. **(B)** Corresponding quantification of evagination resorption dynamics after stretch. **(C)** Time constants resulting from the exponential fitting of the curves in panel (B). Statistical significance was assessed through one-way ANOVA. N=21, 19 and 19 cells from 4 independent experiments. **(D)** Time course images of PM evaginations after stretch on IRSp53^-/-^ MEF expressing either a constitutively active (G12V) form of Rac1 or EGFP-IRSp53-FL. PM was tagged with either GFP for Rac1-G12V and Empty cells or with mCherry for the EGFP-IRSp53-FL transfected cells. **(E)** Corresponding quantification of evagination resorption dynamics after stretch. **(F)** Time constants resulting from the exponential fitting of the curves in panel (E). Statistical significance was assessed through Kruskal-Wallis test. N= at least 19, 16 and 16 cells from 3 independent experiments. For all images scale bars are 5μm.

**Suppl. Fig. 6:**
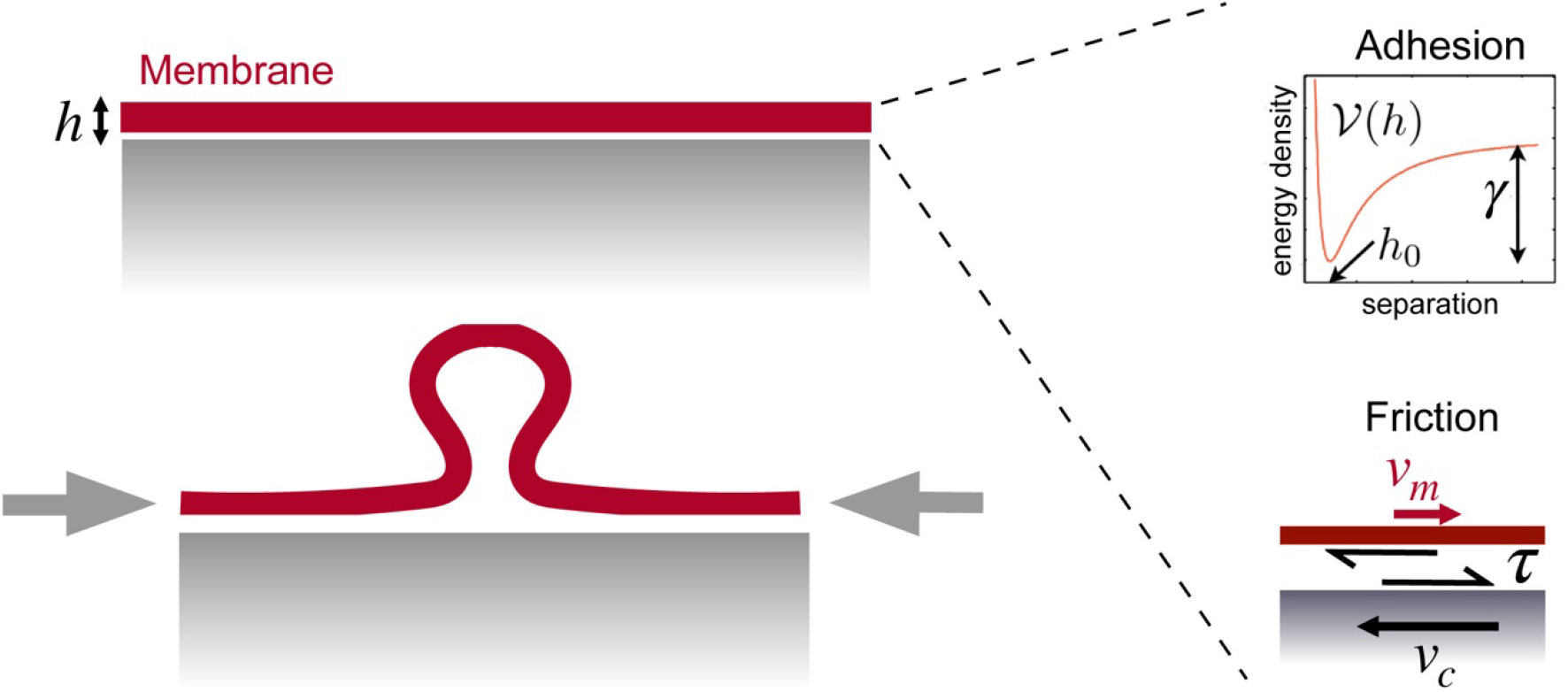
Considerations for the model. Schematic of the interaction between the membrane and the 2D underlying cortex, separated by a distance h before evaginations form. The interaction is modelled through an adhesion potential ν(h) with a minimum at separation h_0_, with adhesion tension γ and a tangential fictional traction τ in the adhered part of the membrane proportional to the slippage velocity *v_m_* — *v_c_*.

## Supplementary Videos

**Supplementary video 1** Time lapse of an NHDF cell labelled with GFP-membrane before, during, and after stretch application. Images on the right side show a magnification of the areas marked in red on the left side.

**Supplementary video 2** Time lapse of an NHDF cell labelled with Actin Chromobody-GFP (ACG) and mCherry-membrane, before, during, and after stretch application. Images on the right side show a magnification of the areas marked in red on the left side.

**Supplementary video 3** Time lapse of an NHDF cell labelled with mEmerald-Ezrin and mCherry-membrane, before, during, and after stretch application. Images on the right side show a magnification of the areas marked in red on the left side.

**Supplementary video 4** Time lapse of a stable NHDF cell line expressing IRSp53 shRNA, labelled with GFP-membrane, before, during, and after stretch application. Images on the right side show a magnification of the areas marked in red on the left side.

**Supplementary video 5** Time lapse of a stable NHDF cell line expressing control Non-Targeting shRNA, labelled with GFP-membrane, before, during, and after stretch application. Images on the right side show a magnification of the areas marked in red on the left side.

**Supplementary video 6** Time lapse of an IRSp53^-/-^ MEF cell, labelled with GFP-membrane before, during, and after stretch application. Images on the right side show a magnification of the areas marked in red on the left side.

**Supplementary video 7** Time lapse of an IRSp53^-/-R^ MEF cell, labelled with GFP-membrane before, during, and after stretch application. Images on the right side show a magnification of the areas marked in red on the left side.

**Supplementary video 8** Time lapse of an IRSp53^-/-^ MEF cell reconstituted with EGFP-IRSp53-FL and labelled with mCherry-membrane before, during, and after stretch application. Images on the right side show a magnification of the areas marked in red on the left side.

**Supplementary video 9** Time lapse of an IRSp53^-/-^ MEF cell, labelled with Actin Chromobody-GFP (ACG) and mCherry-membrane before, during, and after stretch application. Images on the right side show a magnification of the areas marked in red on the left side.

**Supplementary video 10** Time lapse of an IRSp53^-/-R^ MEF cell, labelled with Actin Chromobody-GFP (ACG) and mCherry-membrane before, during, and after stretch application. Images on the right side show a magnification of the areas marked in red on the left side.

**Supplementary video 11** Time lapse of an IRSp53^-/-R^ MEF cell, labelled with pYFP-membrane. Cell is submitted to hypotonic treatment; the medium is subsequently restored to the initial isotonic condition Images on the right side show a magnification of the areas marked in red on the left side.

**Supplementary video 12** Time lapse of an IRSp53^-/-^ MEF cell, labelled with pYFP-membrane. Cell is submitted to hypotonic treatment; the medium is subsequently restored to the initial isotonic condition. Images on the right side show a magnification of the areas marked in red on the left side.

**Supplementary video 13** Time lapse of an IRSp53^-/-^ MEF cell, reconstituted with EGFP-IRSp53-4KE and labelled with mCherry-membrane, before, during, and after stretch application. Images on the right side show a magnification of the areas marked in red on the left side.

**Supplementary video 14** Time lapse of an IRSp53^-/-^ MEF cell, reconstituted with EGFP-IRSp53-I268N and labelled with mCherry-membrane, before, during, and after stretch application. Images on the right side show a magnification of the areas marked in red on the left side.

**Supplementary video 15** Time lapse of an IRSp53^-/-^ MEF cell, reconstituted with EGFP-IRSp53-ΔIBAR and labelled with mCherry-membrane, before, during, and after stretch application. Images on the right side show a magnification of the areas marked in red on the left side.

**Supplementary video 16** Time lapse of an IRSp53^-/-^ MEF cell, reconstituted with EGFP-IRSp53-I408P and labelled with mCherry-membrane, before, during, and after stretch application. Images on the right side show a magnification of the areas marked in red on the left side.

**Supplementary video 17** Time lapse of an IRSp53^-/-^ MEF cell, reconstituted with EGFP-IRSp53-W413G and labelled with mCherry-membrane, before, during, and after stretch application. Images on the right side show a magnification of the areas marked in red on the left side.

**Supplementary video 18** Time lapse of an IRSp53^-/-^ MEF cell, reconstituted with EGFP-IBAR and labelled with mCherry-membrane, before, during, and after stretch application. Images on the right side show a magnification of the areas marked in red on the left side.

**Supplementary video 19** Time lapse of an IRSp53^-/-R^ MEF cell treated with 10μM Wiskostatin and labelled with EGFP-membrane, before, during, and after stretch application. Images on the right side show a magnification of the areas marked in red on the left side.

**Supplementary video 20** Time lapse of an IRSp53^-/-R^ MEF cell treated with 15μM SMIFH2 and labelled with EGFP-membrane, before, during, and after stretch application. Images on the right side show a magnification of the areas marked in red on the left side.

**Supplementary video 21** Time lapse of an IRSp53^-/-R^ MEF cell treated with 10μM para-nitroblebbistatin and labelled with EGFP-membrane, before, during, and after stretch application. Images on the right side show a magnification of the areas marked in red on the left side.

**Supplementary video 22** Time lapse of an IRSp53^-/-R^ MEF cell treated with 25μM CK-666 and labelled with EGFP-membrane, before, during, and after stretch application. Images on the right side show a magnification of the areas marked in red on the left side.

